# Molecular Grammar of Microtubule-Wetting Condensates

**DOI:** 10.1101/2025.10.20.683522

**Authors:** Aswin Vinod Muthachikavil, Christoph Allolio, Thomas D. Kühne

## Abstract

Microtubule condensate interactions are fundamental for cell division, vesicle transport and cellular locomotion. Accordingly, they represent a large number of attractive drug targets. Due to the size of microtubuli and the slow timescale of condensate structural relaxation, there has not been a systematic investigation at the molecular level as to what binding patterns (molecular grammar) enable condensate binding to microtubuli. We provide a protocol that is able to predict whether any given disordered protein sequence will bind to microtubuli. This protocol is suitable for compound screening. Our pattern analysis allows us to establish two categories of strongly interacting subsequences that enable binding to microtubuli: positively charged hydrophobic clusters and alternating charge sequences. Their overall optimal balance is analyzed and preferential regions of interaction on microtubuli are identified and validated with known experimental results. Our results enable rapid prototyping of proteins that target the microtubule surface, i.e. they predict whether unstructured proteins will wet the microtubule interface.

## 1 Introduction

Microtubuli (MT) are the strongest generators of directed mechanical force in the cell, forming “the backbone” of the cytoskeleton [1–3]. Apart from maintaining cellular structure, internal transport processes and locomotion, they also are key actors in cell division: They exert the mechanical force necessary for the separation of chromosomes via the spindle apparatus [4–7]. In animal cells, microtubule nucleation and organization is mediated by the centrosome. Examining centrosome assembly [8, 9], as well as microtubular growth, has resulted in key contributions by the group of Anthony Hyman and many others [10–17], revealing the importance of liquid-liquid phase separation as an organizing principle of the cell. MT-associated proteins, such as Tau, often contain intrinsically disordered regions (IDR), which can undergo phase separation into biomolecular condensates from the surrounding cytosol [18–21]. Their preferential interactions with other proteins then facilitate nucleation, for example, by increasing local concentration and restricting diffusion. As long as the condensate scaffold is a liquid, its interaction with protein interfaces can also be understood as a form of wetting. The biophysical implications of condensate wetting are a very active area of current research [22, 23], as interactions mediated by condensate surface tension are known to mediate centrosome self-assembly, MT lattice formation [24], and branch nucleation [25] that influence force generation by blocking motor proteins [26, 27]. Due to their fundamental importance in cell division, MTs have been a drug target, in particular in oncology [28] and it has even been speculated that they are responsible for the emergence of consciousness [29].

On a more sober note, a detailed molecular understanding of microtubule-condensate interactions will provide a host of new possibilities for the manipulation of fundamental cell processes, including the division of cancer cells [30] and the cure of condensate-specific pathologies, such as tauopathies [31]. Atomistic molecular simulations of condensates on MT remain inaccessible due to the large size of the systems involved. Recent coarse-grained molecular dynamics (MD) simulations [25] have never been shown to exhibit selectivity to predict MT-condensate binding or characterize the MT-condensate interface in sufficient detail to characterize binding in a predictive way. Similarly, machine learning-based models [11, 12, 32] to predict condensate formation and partitioning are not well suited to the detailed interactions involved in IDR-MT wetting, and experimental data remain sparse. Therefore, we set out to establish the molecular selectivity of the recently developed CALVADOS forcefield [33, 34] for MT-condensate binding. Analyzing the resulting binding interface and patterns then allows us to generate a molecular grammar of MT-wetting by intrinsically disordered protein (IDP) condensates.

## 2 Results and Discussion

Determining the molecular basis of the selectivity of MT-wetting by condensates requires a careful validation of the underlying physical model. Hence, we examine the best-known condensate-forming IDPs that bind to MT together with a chemically diverse set of IDPs, which are known to not bind to MTs in experiment. The full sequences and origins of the IDRs/IDPs are given in Section S1 of the Supporting Information.

The IDRs of Tau, LEM2 and TPX2 are able to form condensates that wet the surface of MTs [16, 17, 26, 35, 36]: The longest human Tau isoform, Tau 2N4R, is an IDP that increases the stability of the MT and regulates the interaction of MT-associated proteins (MAPs) [27, 37–42]. Tau35 is a 35 kDa fragment (from E187 to L441) in the C-terminal region of Tau 2N4R, which contains MT-binding domains and is reported to have a higher propensity to aggregate when compared to the full-length Tau [43]. TPX2 (targeting protein for Xklp2) is a protein that stimulates MT branching nucleation. It also plays an important role during apoptosis and is required for chromatin and/or kinetochore-dependent nucleation of microtubuli [17, 35, 36]. As a last example, the proline-arginine-rich low-complexity domain (LCD) at the N-terminal domain of the LEM2 (LEM2-LCD) condenses on the surface of the MT, which is crucial for its role in the nuclear-envelope reformation [16].

On the other hand, the IDRs of FUS, LAF1, and DDX4 are examples of IDPs that form condensates, but have been experimentally confirmed not wet the surface of MTs [44]. However, a combined FUS-Tau2N4R protein was reported to wet the MT surface [45], highlighting the role of binding motifs.

We first establish that all the IDPs examined do indeed form condensates in MD simulations. Condensate formation of the considered IDRs was tested using slab geometries [46–48], as shown in Fig. 1A. The slab simulation protocol provides the densities of the two phases, from the density profile along the interface normal, as shown in Fig. 1B. By varying the temperature, we obtained phase diagrams, with one example shown in Fig. 1C. The phase diagrams of the other IDRs studied in this work, and their estimated upper critical solution temperatures (UCST) are reported in Fig. S2 in the Supporting Information. We observe that the UCST (T_*c*_) of the IDRs varies widely in our simulations, from ∼215 K for Tau35 to ∼323 K for LAF1-RGG, respectively. The densities (mM) of the low-density phases of IDRs at their corresponding 0.92T_*c*_ are shown in Fig. 1D. They do not exhibit any clear correlation with the UCST, showing the importance of actually computing the phase diagrams. In order to have a comparable physical state of the condensate, we chose 0.92T_*c*_ as the reference simulation temperature. Working close to the UCST also ensures fast equilibration. The lowest T_*c*_ belongs to Tau35, a full Tau sequence has even been used as a negative control in the creation of the underlying coarse-grained force field [34, 49]. Regardless, condensate formation occurs both in simulation, as well as in experiment. It is, however, quite difficult to achieve phase separation in absence of nucleation by MTs, as it requires crowders and specific conditions [20, 21].

**Fig. 1.**
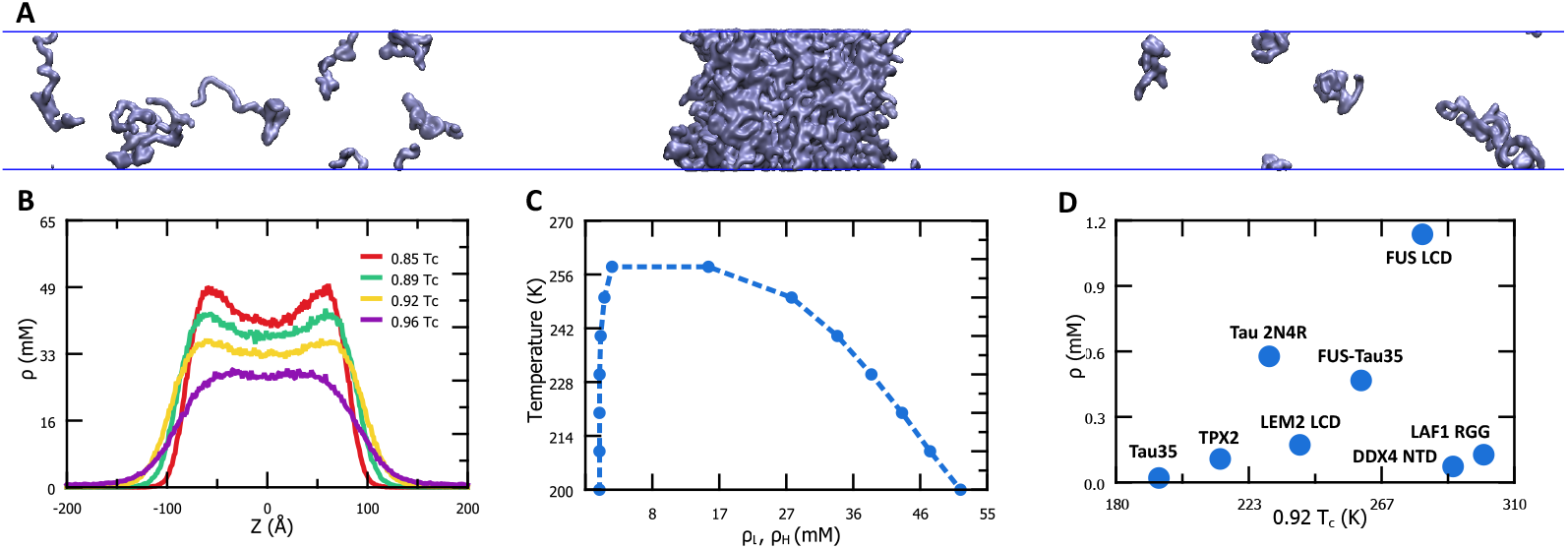
**A**. Co-existence of a high-density and a low-density phase in ‘slab simulations’ of the LEM2-LCD. The normal to the interface of the two phases is along the *z*-axis. **B**. Density profile normal to the interface between high-density and low-density phases. **C**. The phase diagram of the LEM2-LCD condensate, estimated from the density profile. **D**. Densities of the low-density phases (in mM) at 0.92 T_*c*_, where T_*c*_ is the critical temperature.

### 2.1 Microtubule-wetting

The MT-wetting behavior of IDR-condensates was simulated using a restrained, fully periodic MT (see section S3 of the Supporting Information). To better equilibrate the MT interface of bound proteins, we used ‘escape simulations’ at the reference temperature of 0.92T_*c*_. Initially, a biasing force was applied to pull the IDP chains to the surface of the MT. The biasing force was removed for sampling and we monitored whether the condensate remained spread on the MT surface. Condensates that wet MTs remain spread on its surface, whereas those that do not reduce contact with it over time: Fig. 2A shows representative snapshots of LEM2 LCD (wets the MT surface) and FUS-LCD (does not wet the MT surface) from our simulations. Fig. 2B presents the percentage of IDR atoms bound to the surface of the MT, relative to the number bound at time *t* = 0. For IDRs that do not wet the surface, contact with the MT surface is spontaneously reduced over time. A movie visualising the trajectories of all the IDRs used in the study is available in the Supporting Information. We find that our simulations are in agreement with experimental data for all examined IDPs: Proteins known to bind MTs are found to wet the MT surface, and proteins known not to bind are withdrawing from the MT interface. This raises the question, whether this result is trivial in the sense that MT binding proteins sequences have obvious features that are reproduced by the force field: IDRs forming condensates on the MT surface are more hydrophobic, as shown by their significantly higher Uversky hydropathy [50] (a hydrophobicity scale) compared to the others. Electrostatic interactions also play a role in determining the ability of IDR condensates to wet the surface of the MT. Fig. 2D shows the composition of the IDRs. The ones that wet the overall negatively charged MT surface predictably have a higher fraction of basic amino acids (Arginine and Lysine) compared to acidic amino acids (Aspartic acid and Glutamic acid). The non-binding LAF1-RGG, whose repeating RGG (arginine-glycine) units contribute to a higher fraction of positively charged residues, is an exception. It appears, that a specific balance between charge and hydrophobicity is required. Fig. 2C provides an Uversky plot [50] of the simulated IDPs, which plots these quantities against each other. The boundary line (shown in red in Fig. 2c) is a well-known heuristic used to predict whether a protein is folded or disordered. The elevated net charge and hydrophobicity of the MT-wetting IDRs puts them close to the boundary line, indicating the possibility of protein refolding on the MTs. There is experimental evidence for this to occur: The TPX2 *α*_5_-*α*_7_ domain in *Xenopus laevis* is reported to have a disordered state in solution and an ordered structure when bound to MTs [51], as has a TAU fragment [52]. The very strongly coarse-grained CALVADOS model is intrinsically unable to predict such structures. The fact that it is able to predict MT binding regardless of this limitation is a remarkable finding. Hybridizing proteins will scramble their global hydrophathy profiles, but keep binding domains intact, such as is the case for Tau35-FUS. Such test cases enable us to understand what the model provides beyond the information included in the Uversky plot.

**Fig. 2.**
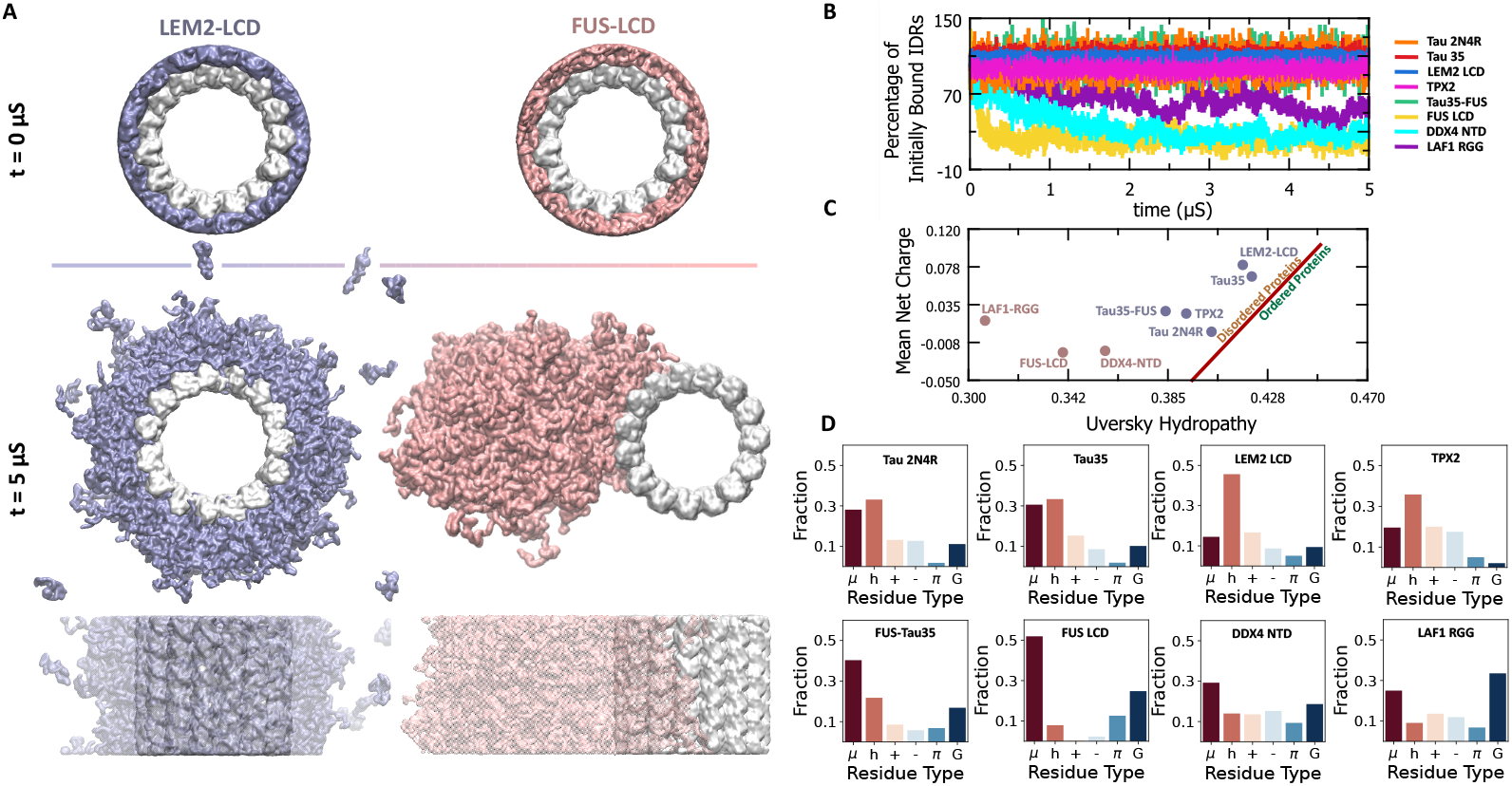
**A**. Selectivity in MT-wetting activity of IDRs exhibited in MD simulations of LEM2-LCD and FUS-LCD on the surface of MT. The MT is shown in white. LEM2-LCD and FUS-LCD condensates are shown in blue and pink, respectively. LEM2-LCD and FUS-LCD both form homogeneous condensates, but only LEM2-LCD wets the surface of MT, while the FUS-LCD condensate does not. Movies visualising 10*µ*S trajectories of all the IDRs studied in this work are available in the Supporting Information. **B**. Number of chains bound to the surface of the MT (as a percentage of the initially bound chains) as a function of time. **C**. The mean net charge vs the Uversky Hydropathy for IDRs used in the study. **D**. The composition of IDRs. The fraction of polar (*µ*), hydrophobic (h), positively charged (+), negatively charged (-), aromatic (*π*), and Glycine (G) are shown.

### 2.2 Cohesion in the condensates

The binding of condensates to MT necessitates the breaking of interactions within the condensate in order to create an interface. The internal interactions and structure of a condensate may already hint at whether it will bind to MT or not. To elucidate this, we performed simulations of the condensates in bulk and quantified interaction probabilities based on a residue-residue interaction energy cutoff of −0.1 kT (see section S2.1 in the Supporting Information).

The dominant pairwise interactions that stabilize the condensate are summarized in residue-residue contact maps (see Fig. S6-S13 of the Supporting Information). Yet, residues in the IDR chain do not interact with each other in a statistically independent manner. We identified ‘blocks’ of residues that tend to cooperate while forming contacts, based on the cooperativity of the contact formation of individual residues within them. Residues with contact probabilities greater than the top *n*^*th*^ percentile are considered ‘active’, where the value of n is chosen to globally maximize the correlations within blocks: A block is defined on the IDP sequence as a set of consecutive, active residues. Additionally, active residues separated by a maximum of one non-active residue can belong to the same block. In order to estimate the (IDP specific) cutoff for block formation, we use a negative Kullback-Leibler divergence to estimate mutual information by

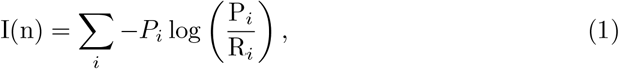

where P_*i*_ is the joint probability that all the ‘active residues’ in a block *i* make contacts simultaneously with a common chain. R_*i*_ is the probability that the constituents of the block *i* form contacts independently. Constituents of a block can either be single residues or sub-blocks found at a higher cutoff. This can be interpreted as a truncated version of the total correlation [53], restricted to independent contacts (of sub-blocks) and the full-size block. Blocks are built up iteratively from those found at the previous cut-off (See Methods for detailed explanation). *I*(*n*) thereby provides a measure of the cooperativity of contacts within residues in the blocks of an IDR chain: Negative values of I(n) indicate a higher cooperativity among the residues of a block to form contacts. The value points to how much information the actual contact pattern provides compared to what we would expect from an independent binding by its constituents. The clustering cutoff for each protein was set to the contact probability percentile cutoff that captures the highest mutual information content in the sense of Eq. 1, compared to the individual constituents of the respective block (See Fig. 3A and section S4.1.1 in the Supporting Information). Fig. 3B shows the contact blocks in the Tau 2N4R chain. The residues that are a part of a block are labeled together. Fig. S15 shows the blocks identified in all the IDRs used in the study by this methodology.

**Fig. 3.**
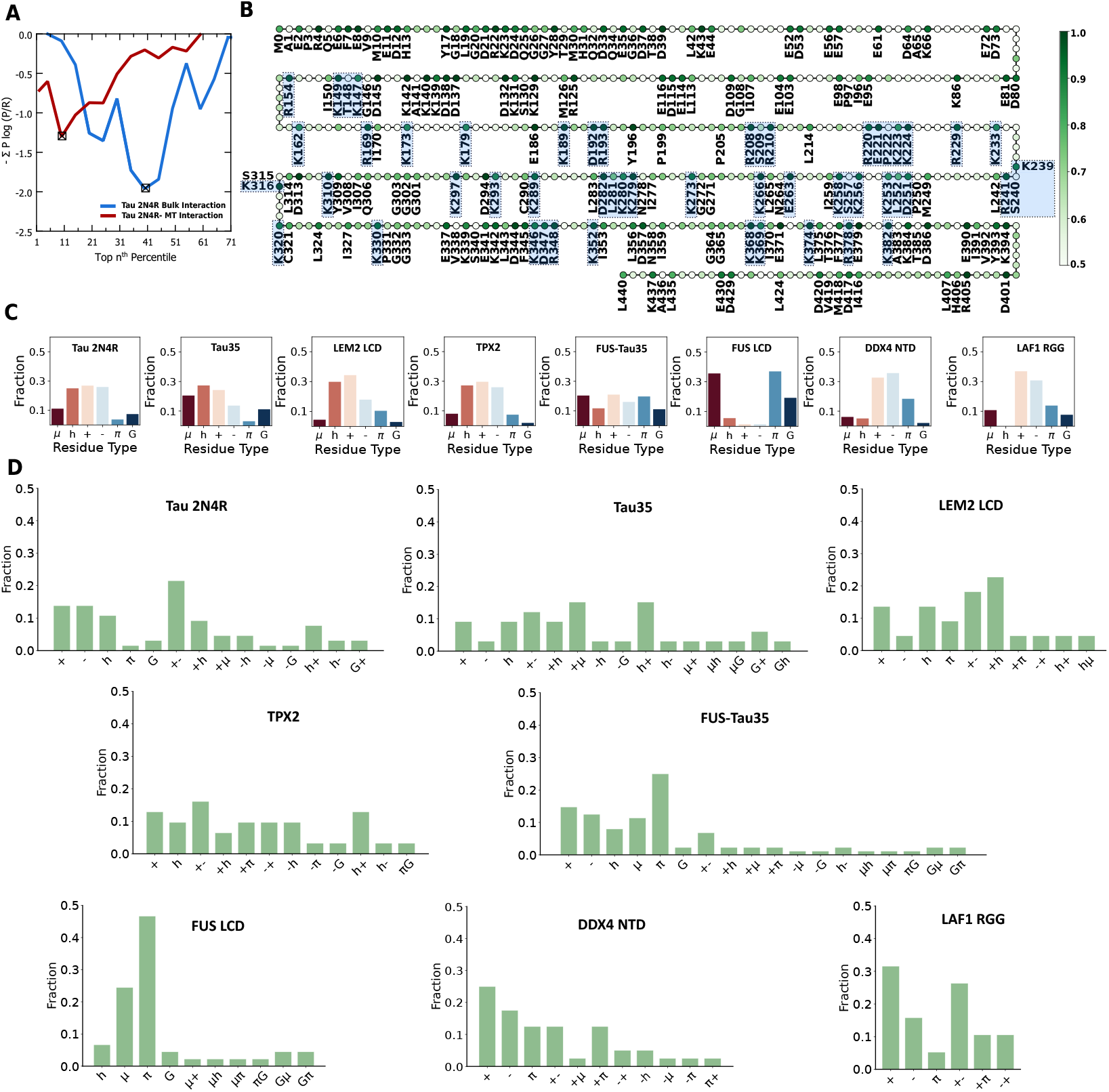
**A**. The randomness of contact-formation of active residues of blocks (Tau 2N4R) plotted against the condition for selecting active residues. If a residue has a probability of forming a contact in the top *n*^*th*^ percentile, it is identified as an active residue. The blue line represents the contact formation within the condensate bulk, and the red line represents the contact between IDR and the MT surface. A block is determined based on selecting active residues that minimize the randomness of contact formation. **B**. Visual representation of contact blocks in Tau 2N4R. Each circle represents a residue. The residues that are labeled in the chain represent the blocks identified in the IDR chain. The residues highlighted in blue are the blocks that interact with the MT surface. The colour of the circle represents the probability of the residue to make a contact in the bulk of the condensate at constant pressure. **C**. The average residue type composition of blocks in all IDR chains. Residues are categorised into positive (+), negative (-), polar (*µ*), hydrophobic (h), aromatic (*π*) and glycine (G). **D**. Types of blocks in IDR-condensates, categorized by the dominant and sub-dominant residue type in the block. The first letter shows the dominant residue type, and the second letter shows the sub-dominant residue type in the block.

Essentially, these blocks represent the ‘sticky’ domains of the IDR chains that stabilize their condensates, and hence reveal the nature of interactions essential to their formation. The average compositions of the contact blocks of various IDRs are shown in Fig. 3C. In almost all IDRs, charged amino acids make up a big part of blocks, indicating their significance in the formation of condensates. An exception to this is the FUS LCD, where the blocks are significantly comprised of polar and aromatic residues. We note the enrichment of hydrophobic residues in the contact blocks of the MT-wetting IDRs, compared to the non-wetting IDRs. Furthermore, the neighborhood of the contact blocks is, in general, more hydrophobic for the MT-wetting IDRs (see Fig. S16 of the Supporting Information), compared to that of non-MT-wetting IDRs.

To gain further insight into the nature of interactions, we categorized blocks based on the dominant and subdominant residue types in them. The dominant residue type is the kind of residue that appears most frequently, and the subdominant residue type appears the second most frequently. For example, a block with only positive residues is a labeled positive (+). Blocks with mostly positive residues and hydrophobic residues as the second most common type are labeled positive-hydrophobic (+h). Electrostatic interactions, respectively salt bridges are abundant for all IDPs, but FUS LCD. Alternating charge patterns inside interacting blocks are common in both MT binding (LEM2, TAU) and nonbinding condensates (LAF1 RGG, DDX4 NTD), which is why we distinguish between charged residues. A general similarity is observed between the types of blocks in Tau2N4R, and its smaller Tau35 fragment. In the case of the LEM2-LCD, the majority of its blocks are positive-hydrophobic residues. Among domains without charges, hydrophobic and aromatic residues play the major role.

An interesting case is that of the combined FUS-Tau35. We observe that FUS-Tau35 has contributions from the individual FUS LCD and the Tau35 domains to the block-types. FUS domain is largely stabilised by blocks with polar, aromatic, and hydrophobic residues in them. On the other hand, Tau35 is rich in blocks containing charged residues. The combined FUS-Tau35 protein has blocks heavily enriched with charged residues (similar to Tau35), and blocks that are dominated by hydrophobic, polar and aromatic residues as well. We assume that different fragments of the IDRs contribute in a distinct way towards the stability of the condensate. However, we also note the disappearance of some block types in the combined FUS-Tau35.

A key difference between the MT-wetting and MT-non-wetting IDRs is the absence of +h or h+ blocks in the latter. Notably, not only the hydrophobicity of the blocks, but also of their neighborhood is higher for the IDRs that wet the surface of MT (see Fig. S16 in the Supporting Information). Aromatic contributions are common in DDX4, FUS, and LAF1 RGG, but also in the FUS-Tau35 hybrid. Our analysis provides a complement to pure sequence-based pattern statistics, in that only the major interacting parts of the IDP sequences are considered. This focuses analysis to the most important subsequences. The non-random arrangement of residues in disordered regions inferred using numerical intermixing (NARDINI) provides a good measure for binary patterns of residue types with respect to one another in an IDR sequence [54]. The binary residue-type patterning on the whole IDR sequences, analysed using NARDINI, also indicates an interlaced hydrophobic-basic residue patterning for IDRs that wet the MT surface (see Fig. S18 and associated discussion in the Supporting Information. While it provides more detail on a molecular level, it essentially replicates the insight from the Uversky plot.

### 2.3 Microstructure of MT-Wetting

So far, our results indicate that condensates that wet the MT surface have a distinct molecular grammar, defined by the presence of a high degree of hydrophobicity clustered together with positive charges into interacting blocks. Yet, apart from the presence of these substructures, a diverse set of interactions is found for MT-wetting proteins. Formation of secondary structures in previously disordered regions bound to the surface of MT is known to occur [51]. If the binding to MT was dependent on refolding on the surface, it is unlikely that our simulations would be able to reproduce the selectivity. We, therefore, propose that condensate interactions with the MT surface are, to a certain extent, independent of the protein’s secondary structure. This interpretation of MT-condensate interactions has been proposed previously [16], by declaring MT-condensate interactions “unspecific”, but without molecular evidence.

To clarify the difference, we distinguish sequence *chemical* specificity from secondary *structural* specificity. We use chemical specificity in the sense of specific ion effects, where ions of identical charge have different effects on surface tension. Accordingly, we provide density profiles at the MT interface resolved by the residue type in Fig. 4A. For all IDPs considered, we observe that the condensate interface at the surface of MT has significant hydrophobicity. This is consistent with the finding of a higher Uversky hydropathy and a generally higher hydrophobic composition for these proteins. With the exception of Tau35, which has the highest hydrophobicity, we find an increase in the density of hydrophobic residues at the MT-condensate interface. The density of positive charge also increases, with the exception of LEM2, which has the highest bulk charge. This indicates a preferential interface composition (chemical specificity) of the IDPs binding to MTs, limiting hydrophobicity and charge accumulation. It appears that the ideal composition lies between that of LEM2 and of Tau35. However, local structure ultimately determines binding. The residues at the surface of the MT that interact with the condensates are shown in Fig. 4B. (see the Supporting Information for IDR-MT contact data). A dominant fraction of contacts are between the negatively charged surface residues of the MT and the positively charged residues in the IDPs. The overall charge of the MT surface is slightly negative, but there are also several positively charged residues exposed (see Fig. 4C). We examined the molecular composition of the average neighborhood (within 0.5 nm) of residues on the surface of the MT, as shown in Fig. 4C. Positively charged surface MT residues, on average, have slightly fewer hydrophobic residues at the interface in their environment. However, they have a larger fraction of negatively charged residues in their immediate vicinity, thereby reducing the local net charge. Negatively charged surface residues of the MT, on the other hand, have a significantly larger proportion of hydrophobic residues in their surface neighborhood, and fewer positively charged ones. Accordingly, locally highly negatively charged and hydrophobic neighborhoods exist on the MT surface. These negatively charged neighborhoods form hotspots of interaction with the cationic residues of the IDR (see Fig. 4D). This observation consistently explains the selectivity of positively charged hydrophobic IDRs wetting the surface of the MT. The cationic binding centers on the MT provide an alternative binding mode of alternating charges, which might also benefit from the presence of hydrophobic residues. To better understand these observations, we fitted a random forest model to the binding data (see Supporting Information for details). Fig. 4E shows the partial dependence of an IDR residue (of the probability that it makes a contact with MT) on the presence of positive and hydrophobic residues in the neighborhood. It is not surprising that for positive residues, both the presence of positive and hydrophobic residues in their vicinity enables the formation of a contact with the MT. For negative residues, on average, the presence of positive charges enables their binding onto the surface of the MT. However, the presence of hydrophobic residues in the vicinity does not affect their MT-contact probabilities. This illustrates the two different binding modes of positively charged and negatively charged IDP interacting groups on MTs.

**Fig. 4.**
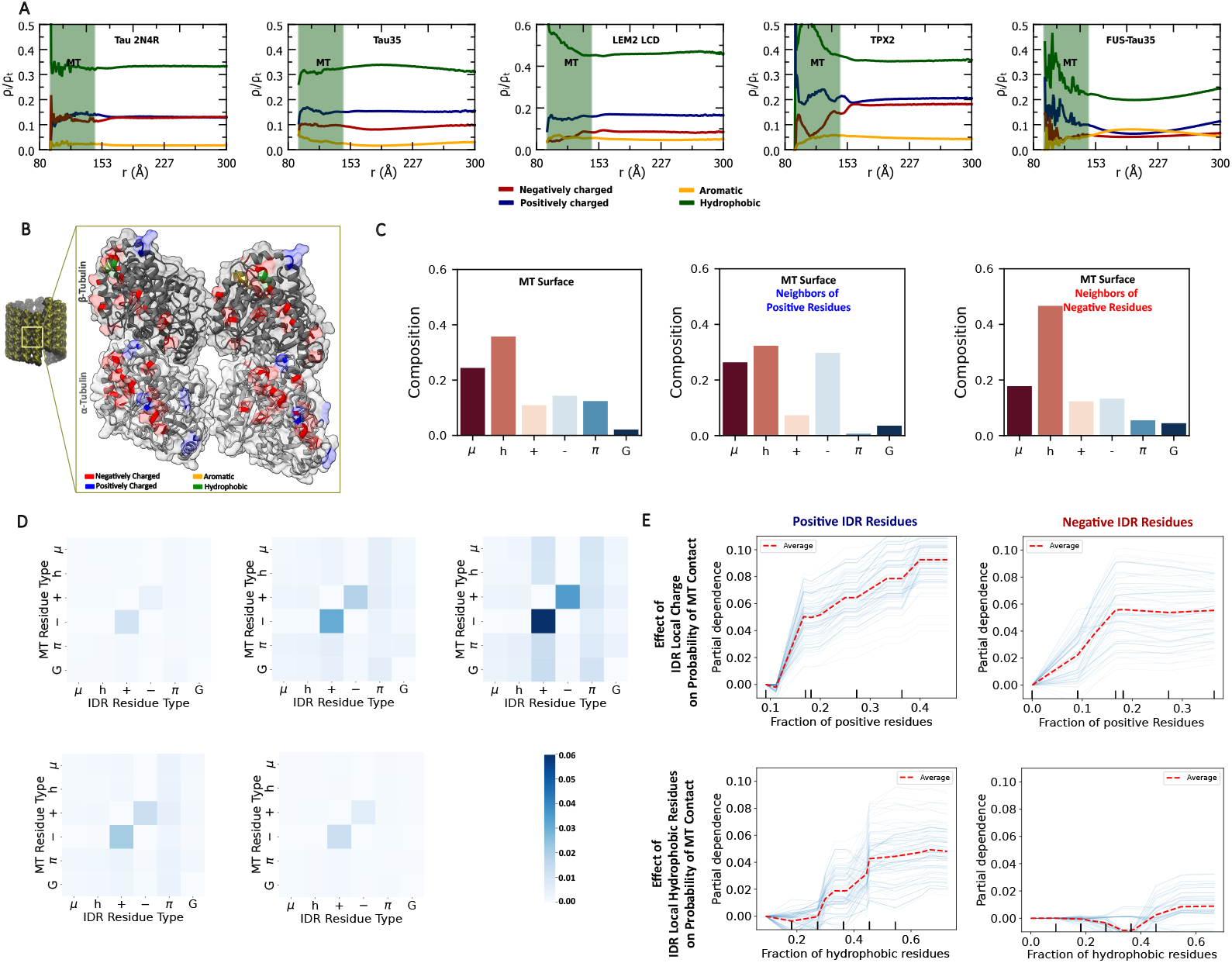
**A**. The radial density profiles of different residue types on the surface of the MT. The shaded green region represents the MT-wall. **B**. MT surface with highlighted residues that form contacts with the IDR residues most frequently. Positive (blue), negative (red), aromatic (orange), and hydrophobic (green) residues are highlighted using different colours. **C**. The surface composition of MT. The average neighborhood of positive, negative residues on MT surface are also shown separately. **D**. Heat map showing the interaction between residues of IDR and MT, based on residue types. **E**. Partial dependence of MT-contact probability of residues estimated by a random-forest regressor model. The red line indicates the average predicted value of the model as a function of a single feature, marginalizing over all other features. The blue lines represent individual predicted responses for a subset of input samples when the target feature is varied, and all other features are held fixed. The partial dependences of (positively and negatively charged residues) on the presence of nearby positive residues and nearby hydrophobic residues are shown separately.

In view of the low resolution of the force field used, it is reasonable to doubt the completeness of this picture. Several published experimental studies characterize condensate binding to microtubuli. In particular for TAU and TPX2 existing studies classify MT binding domains as well as structural data of the binding. We first discuss a detailed TPX2 NMR study on *Xenopus Laevis* TPX2, which includes evidence of refolding of the relevant protein on the MT [51]: The sequence studied by Guo et al. is closely related to the human TPX2 IDR we studied. One difference is that the 477-716 domain of the human TPX2 used in our simulations does not include the so-called Eg5 domain[51], which was however verified to not influence the MT-binding affinity of the protein [51, 55, 56]. Guo et al. studied the C-terminal region of a TPX2 domain, i.e. the minimal active fragment (*α*5-*α*7 with amino acids 477–716 of the full sequence) that binds to MTs and enables branching MT nucleation. They report refolding of this IDP upon binding to MT. Furthermore, TPX2^*α*5*−α*7^ was found to bridge two adjacent heterodimers on the MT. This is in general agreement with our contact analysis, but obviously without the structural specificity of refolding: The most important residues binding to TPX2 lie at the interface between adjacent tubulins on the MT surface. The specific residues of the *α*- and *β*-tubulins that bind to TPX2 in our study are shown in Fig. S17 of the Supporting Information. From our analysis, D394, R400, E412, E413, E431, E432, and D436 of the *α*-tubulins, and E155, R388, R389, E399, D402, E403, M404, E405 and D425 of the *β*-tubulin are the residues at the interface between adjacent tubulins that contact the TPX2 condensate favorably (The residue indices start at 0). We note that, in contrast to the experimental TPX2 NMR findings, the binding residues are mostly charged aminoacids.

TPX2 also contains multiple FKARP motifs, which are reported to interact with MT [55]. The FKARP motif is also conserved in the human TPX2 that was used in this study, and contains a contact block in it that interacts with the MT (block 149-151 shown in Fig. S15 in the Supporting Information). Conserved FKARP and FKAQP (FKALP in human TPX2) motifs of *Xenopus Laevis* TPX2 have been reported to face away from the MT in the refolded structure[51], but simultaneously contacts by the constituting charged aminoacids have been reported in the same study. It is entirely possible that these strong interactions originate from unstructured parts of the condensate and that the reported refolded structure is just a minor binding mode, which happened to be amenable to NMR spectroscopy. Regardless, our results are compatible with the region of preferential binding on the MT in the sense that binding residues are concentrated at the edges of the tubulin molecules. Specific domains have also been reported to play an important role in in MT-IDR interactions[57] of TAU 24NR. From the residue-level contact analysis performed in this work, these domains contain highly active residues that interact with the MT. A 12-residue repeat in the R1 (R241-G272) region (VKSKIGSTENLK) was identified in their work as the best-resolved segment forming critical interactions with the MT. This region in the R1 of Tau 2N4R corresponds to two MT-contact blocks identified in our study (K256-K258 and E263-K266). The NKK residues in the VQIINKK sequence of R2 (K273-G303) are also identified as a strongly MT-interacting domain of Tau. Kellogs et al. [57] also report that TAU 2N4R interacts with the residues at the interface between *α* and *β* tubulins. This is also in agreement with our results because we find that MT residues at the interface of the *α*-(D436,E432,E431,D394), and *β*-(D425,E155,E403,E405,R388) tubulins make contacts with Tau2N4R.

These observations in the literature are in line with the earlier results by Tan et. al. [58], who reported that the lattice of the MT could be important in the stabilization of condensate on its surface. They observed that Tau condenses on taxol-stabilized or native guanosine diphosphate (GDP) MT lattices, but not on MTs stabilized with the non-hydrolyzable GTP analogue GMP-CP.

The full-length Tau-FUS combined IDR has been reported in the literature to wet the MT surface [45]. We combined the Tau35 fragment (which has all four MT-binding repeats [43]) with FUS to check the merged protein’s ability to wet the MT-surface. It was observed that the condensate of this ‘fusion’ protein does wet the surface of the MT in our simulations, in agreement with experimental evidence of hybrid protein binding. Fig. 5A shows the condensate spread on the surface of the MT. As expected, it is the Tau35 residues in the combined FU-Tau35 condensate which we have found to be responsible for favorable interactions with MT, as indicated by the density profile in Fig. 5B. The overall interactions between the FUS LCD residues are stronger than those between Tau35 residues, as indicated by its high critical temperature. Therefore, the FUS residues also bury themselves within the condensate, exposing the Tau35 residues to the condensate-solvent interface to minimize surface tension. Subdomains within an IDR maintain their identity and contribute collectively to the behavior of the entire IDR. This illustrates the limits of statistical analysis of the whole IDR sequence, as opposed to a cluster analysis of local interactions.

**Fig. 5.**
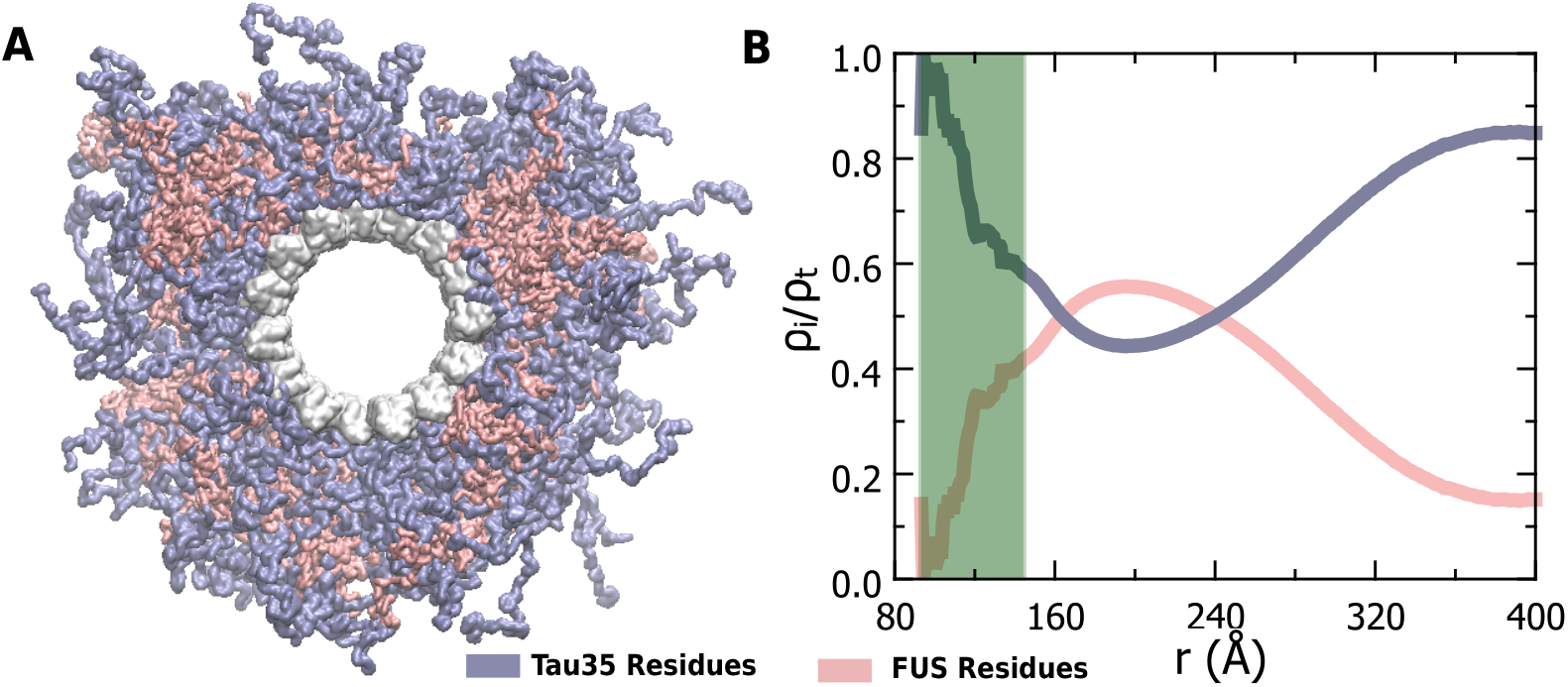
**A**. The condensate of the FUS-Tau35 protein spreads on the surface of the MT surface. The Tau35 (blue) and FUS (red) residues are coloured differently to distinguish their spatial concentration. **B**. The radial density profile of the residues of the Tau35 and FUS domains (*ρ*_*i*_), normalised by the total density (*ρ*_*t*_).

Our coarse-grained simulations have shown to reproduce the selectivity of experimentally observed MT-condensate binding. Moreover, they provide a surprisingly detailed picture of local interactions, enabling us to construct a molecular grammar of MT wetting based on interacting fragments. Two primary interaction modes are observed: (1) electrostatic and hydrophobic, where the condensate carries a positive charge, and (2) alternating charge electrostatic, localized near positively charged microtubule (MT) residues. We also estimated the optimal global hydrophobicity and charge properties and identified the strongly interacting region of the MT interface. Incorporating further data might yield a descriptor that is able to predict binding from sequence alone. Another obvious future step is to use our findings to construct new MT-binding disordered proteins and test our analysis experimentally. We also think our approach might be suitable to explain the observed differences in affinity for curved MT [57], thereby further facilitating protein design and paving the way for targeting MTs in an even more specific manner with a wide range of potential therapeutic applications.

## 3 Methods

### Simulation methods

The CALVADOS3 coarse-grained force-field recently developed by Cao et al. [33] was used to model proteins in the simulations presented in this work. The parameters of the force-field version used are included in the Supplementary Dataset 1. The model is reported to be capable of representing both disordered and multi-domain proteins in simulations. Simulations were performed using OpenMM 8.1.2 [59] in NVT ensemble. Time integration was performed using the Langevin integrator [60] with a friction coefficient of 0.01/ps. Simulations were carried out at a salt concentration of 0.15 mM and a pH of 7.5. A time step of 10 fs was used to perform time integration. Trajectories were written out after every nanosecond. For simulations of bulk-condensate phases, NPT simulations were used with the Monte Carlo barostat [61, 62]. Volume changes were attempted every 25 time steps. In simulations involving MT, all MT atoms were position-restrained by a harmonic spring of spring constant 1000 KJ/mol/nm^2^. A wall potential was applied at the inner surface of the MT cylinder (see section S3 in the Supporting Information).

### Identification of contact blocks

A contact block is a functional domain of an IDR chain, within which the contact-forming (active) residues cooperate while forming contacts. Active residues are residues with a contact-forming probability greater than a cut-off value. It is possible that active residues that are separated from each other by inactive residues can also form contacts co-operatively. To account for this possibility, we allow for a maximum of one inactive residue between two active residues, to be able to be a part of the same contact block.

As mentioned earlier, active residues are those that are decided based on a cut-off probability. We define active residues as the ones that have a probability of forming a contact in the top n^*th*^ percentile among the residues in the IDR chain. As expected, when the cut-off is sufficiently high, fewer residues are selected as active, and therefore result in a large number of blocks with fewer residues. As the value of ‘n’ is increased, a larger number of residues are classified as active residues. These get added to residues, and blocks identified with the previous value of ‘n’ merge, or newer residues get added to existing ones to form larger blocks. The change in average block size and the number of blocks with the value of ‘n’ is shown in Fig. S14 in the Supporting Information file. The joint probability of active residues within a block is the probability that all residues in the block of a chain form a contact simultaneously with a common chain. As expected, the joint probability of active residues decreases with block size (Fig. S14 in the Supporting Information file). Now, we characterize the randomness of active residue contacts in the blocks of an IDR by I(n)=-SP_i_log(P_i_*/*R_i_), as described in equation 1. R_*i*_ is defined as the probabilities of active constituents of the block:

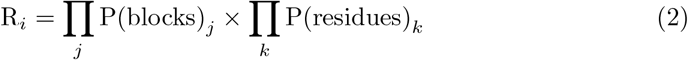

For a given value of *n*, the ‘blocks’ identified for the previous value of *n* get expanded to form new, larger blocks as described earlier (Fig. S14 in Supporting Information shows the average size of blocks with n). For each of the new blocks (*i*), the previous smaller blocks that are completely contained in it, and the additional active residues that make up the new block. The joint probability of these sub-blocks (*P* (*blocks*)_*j*_) and the probability of contact formation of additional active residues (*P* (*residues*)_*k*_) are used to calculate R_*i*_ as described in equation 2. This negative Kullback-Leibler divergence-based metric measures the deviation from random contact formation, with I(n) = 0 indicating perfectly random behavior, I(n) *>* 0 indicating more dispersed contact patterns than random (i.e. anticorrelation), and I(n) *<* 0 indicating more structured organization and higher information content than random expectation.

Fig. S14 in the Supporting Information shows the variation of non-cooperativity in the blocks with *n* for all IDRs in the study. We observe there is an optimal definition for blocks based on the value of ‘n’, for which the active constituents of a block form contacts with most cooperativity. Beyond this value, the randomness of contact formation within a block begins to increase. We define a block based on the value of *n* that corresponds to the minimum of *I*(*n*).

While identifying the dominant and subdominant residue types within a block, ties between residue types were resolved using a residue-type hierarchy based on interaction strength. Electrostatic residues were assigned the highest priority. The remaining residue types were ranked according to their average stickiness parameter (*λ*) values in the force field. The resulting hierarchy used was aromatic (*π*) *>* Glycine (G) *>* hydrophobic (h) *>* polar (*µ*) residues.

## Authors’ Contributions

AV conducted research. CA designed research. AV and CA performed analysis and wrote the paper. CA supervised the project, TK initiated research. TK revised the paper and contributed to the interpretation of the results.

## Acknowledgements

This work has been supported by Charles University Research Centre program No. UNCE/24/SCI/005.

## References

[1] Laan, L., Husson, J., Munteanu, E.L., Kerssemakers, J.W., Dogterom, M.: Force-Generation and Dynamic Instability of Microtubule Bundles. Proc. Natl. Acad. Sci. USA 105(26), 8920–8925 (2008) 10.1073/pnas.0710311105

[2] Mitchison, T., Kirschner, M.: Dynamic Instability of Microtubule Growth. Nature 312(5991), 237–242 (1984) 10.1038/312237a0

[3] Dogterom, M., Kerssemakers, J.W., Romet-Lemonne, G., Janson, M.E.: Force Generation by Dynamic Microtubules. Curr. Opin. Cell Biol. 17(1), 67–74 (2005) 10.1016/j.ceb.2004.12.011

[4] Joglekar, A.P., Bloom, K.S., Salmon, E.: Mechanisms of Force Generation by End-on Kinetochore-Microtubule Attachments. Curr. Opin. Cell Biol. 22(1), 57–67 (2010) 10.1016/j.ceb.2009.12.010

[5] Kent, I.A., Lele, T.P.: Microtubule-Based Force Generation. Wiley Interdiscip. Rev.: Nanomed. Nanobiotechnol. 9(3), 1428 (2017) 10.1002/wnan.1428

[6] Chu, L.-Y., Stedman, D., Gannon, J., Cox, S., Pobegalov, G., Molodtsov, M.I.: Force-Transducing Molecular Ensembles at Growing Microtubule Tips Control Mitotic Spindle Size. Nat. Commun. 15(1), 9865 (2024) 10.1038/s41467-024-54123-2

[7] Elting, M.W., Suresh, P., Dumont, S.: The Spindle: Integrating Architecture and Mechanics across Scales. Trends Cell Biol. 28(11), 896–910 (2018) 10.1016/j.tcb.2018.07.003

[8] Heald, R., Tournebize, R., Blank, T., Sandaltzopoulos, R., Becker, P., Hyman, A., Karsenti, E.: Self-Organization of Microtubules into Bipolar Spindles Around Artificial Chromosomes in Xenopus Egg Extracts. Nature 382(6590), 420–425 (1996) 10.1038/382420a0

[9] Woodruff, J.B., Ferreira Gomes, B., Widlund, P.O., Mahamid, J., Honigmann, A., Hyman, A.A.: The Centrosome is a Selective Condensate that Nucleates Microtubules by Concentrating Tubulin. Cell 169, 1066–1077 (2017) 10.1016/j.cell.2017.05.028

[10] Brangwynne, C.P., Eckmann, C.R., Courson, D.S., Rybarska, A., Hoege, C., Gharakhani, J., Juülicher, F., Hyman, A.A.: Germline P Granules are Liquid Droplets that Localize by Controlled Dissolution/Condensation. Science 324(5935), 1729–1732 (2009) 10.1126/science.1172046

[11] Kilgore, H.R., Mikhael, P.G., Overholt, K.J., Boija, A., Hannett, N.M., Van Dongen, C., Lee, T.I., Chang, Y.-T., Barzilay, R., Young, R.A.: Distinct Chemical Environments in Biomolecular Condensates. Nat. Chem. Biol. 20(3), 291–301 (2024) 10.1038/s41589-023-01432-0

[12] Saar, K.L., Scrutton, R.M., Bloznelyte, K., Morgunov, A.S., Good, L.L., Lee, A.A., Teichmann, S.A., Knowles, T.P.J.: Protein Condensate Atlas from Predictive Models of Heteromolecular Condensate Composition. Nat. Commun. 15(1), 5418 (2024) 10.1038/s41467-024-48496-7

[13] Schneider, M.W.G., Gibson, B.A., Otsuka, S., Spicer, M.F.D., Petrovic, M., Blaukopf, C., Langer, C.C.H., Batty, P., Nagaraju, T., Doolittle, L.K., Rosen, M.K., Gerlich, D.W.: A Mitotic Chromatin Phase Transition Prevents Perforation by Microtubules. Nature 609(7925), 183–190 (2022) 10.1038/s41586-022-05027-y

[14] Alshareedah, I., Borcherds, W.M., Cohen, S.R., Singh, A., Posey, A.E., Farag, M., Bremer, A., Strout, G.W., Tomares, D.T., Pappu, R.V., Mittag, T., Banerjee, P.R.: Sequence-specific Interactions Determine Viscoelasticity and Ageing Dynamics of Protein Condensates. Nat Phys. 20(9), 1482–1491 (2024) 10.1038/s41567-024-02558-1

[15] Banani, S.F., Lee, H.O., Hyman, A.A., Rosen, M.K.: Biomolecular Condensates: Organizers of Cellular Biochemistry. Nat. Rev. Mol. Cell Biol. 18(5), 285–298 (2017) 10.1038/nrm.2017.7

[16] Appen, A., LaJoie, D., Johnson, I.E., Trnka, M.J., Pick, S.M., Burlingame, A.L., Ullman, K.S., Frost, A.: LEM2 Phase Separation Promotes ESCRT-mediated Nuclear Envelope Reformation. Nature 582(7810), 115–118 (2020) 10.1038/s41586-020-2232-x

[17] Bird, A.W., Hyman, A.A.: Building a Spindle of the Correct Length in Human Cells Requires the Interaction between TPX2 and Aurora A. J. Cell Biol. 182(2), 289–300 (2008) 10.1083/jcb.200802005

[18] Islam, M., Shen, F., Regmi, D., Petersen, K., Karim, M.R.U., Du, D.: Tau Liquid– Liquid Phase Separation: at the Crossroads of Tau Physiology and Tauopathy. J. Cell. Physiol. 239(6), 30853 (2024) 10.1002/jcp.30853

[19] Boyko, S., Surewicz, K., Surewicz, W.K.: Regulatory Mechanisms of Tau Protein Fibrillation under the Conditions of Liquid–Liquid Phase Separation. Proc. Natl. Acad. Sci. USA 117(50), 31882–31890 (2020) 10.1073/pnas.2012460117

[20] Kanaan, N.M., Hamel, C., Grabinski, T., Combs, B.: Liquid-Liquid Phase Separation Induces Pathogenic Tau Conformations in Vitro. Nat. Commun. 11(1), 2809 (2020) 10.1038/s41467-020-16580-3

[21] Ambadipudi, S., Biernat, J., Riedel, D., Mandelkow, E., Zweckstetter, M.: Liquid– Liquid Phase Separation of the Microtubule-Binding Repeats of the Alzheimer-Related Protein Tau. Nat. Commun. 8(1), 275 (2017) 10.1038/s41467-017-00480-0

[22] Gouveia, B., Kim, Y., Shaevitz, J.W., Petry, S., Stone, H.A., Brangwynne, C.P.: Capillary forces generated by biomolecular condensates. Nature 609(7926), 255– 264 (2022) 10.1038/s41586-022-05138-6

[23] Wang, Y., Li, S., Mokbel, M., May, A.I., Liang, Z., Zeng, Y., Wang, W., Zhang, H., Yu, F., Sporbeck, K., Jiang, L., Aland, S., Agudo-Canalejo, J., Knorr, R.L., Fang, X.: Biomolecular condensates mediate bending and scission of endosome membranes. Nature 634(8036), 1204–1210 (2024) 10.1038/s41586-024-07990-0

[24] Siahaan, V., Tan, R., Humhalova, T., Libusova, L., Lacey, S.E., Tan, T., Dacy, M., Ori-McKenney, K.M., McKenney, R.J., Braun, M., Lansky, Z.: Microtubule lattice spacing governs cohesive envelope formation of tau family proteins. Nat. Chem. Biol. 18(11), 1224–1235 (2022) 10.1038/s41589-022-01096-2

[25] Biswas, S., Grover, R., Reuther, C., Poojari, C.S., Shaebani, R., Nandakumar, S., Gruünewald, M., Zablotsky, A., Hub, J.S., Diez, S., John, K., Schaedel, L.: Tau accelerates tubulin exchange in the microtubule lattice. Nat. Phys. (2025) 10.1038/s41567-025-03003-7

[26] Siahaan, V., Krattenmacher, J., Hyman, A.A., Diez, S., Hernández-Vega, A., Lansky, Z., Braun, M.: Kinetically Distinct Phases of Tau on Microtubules Regulate Kinesin Motors and Severing Enzymes. Nat. Cell Biol. 21(9), 1086–1092 (2019) 10.1038/s41556-019-0374-6

[27] Chaudhary, A.R., Berger, F., Berger, C.L., Hendricks, A.G.: Tau Directs Intracellular Trafficking by Regulating the Forces Exerted by Kinesin and Dynein Teams. Traffic 19(2), 111–121 (2018) 10.1111/tra.12537

[28] Jordan, M.A., Wilson, L.: Microtubules as a target for anticancer drugs. Nat. Rev. Cancer 4(4), 253–265 (2004) 10.1038/nrc1317

[29] Hameroff, S., Penrose, R.: Orchestrated reduction of quantum coherence in brain microtubules: A model for consciousness. Math. Comput. Simul. 40(3), 453–480 (1996) 10.1016/0378-4754(96)80476-9

[30] Boija, A., Klein, I.A., Young, R.A.: Biomolecular condensates and cancer. Cancer Cell 39(2), 174–192 (2021) 10.1016/j.ccell.2020.12.003

[31] Orr, M.E., Sullivan, A.C., Frost, B.: A brief overview of tauopathy: Causes, consequences, and therapeutic strategies. Trends Pharmacol. Sci. 38(7), 637–648 (2017) 10.1016/j.tips.2017.03.011

[32] Mierlo, G., Jansen, J.R.G., Wang, J., Poser, I., Heeringen, S.J., Vermeulen, M.: Predicting protein condensate formation using machine learning. Cell Rep. (2021) 10.1016/j.celrep.2021.108705

[33] Cao, F., Buülow, S., Tesei, G., Lindorff-Larsen, K.: A Coarse-grained Model for Disordered and Multi-Domain Proteins. Protein Sci. 33(11), 5172 (2024) 10.1002/pro.5172

[34] Tesei, G., Schulze, T.K., Crehuet, R., Lindorff-Larsen, K.: Accurate Model of Liquid–Liquid Phase Behavior of Intrinsically Disordered Proteins from Optimization of Single-Chain Properties. Proc. Natl. Acad. Sci. USA 118(44), 2111696118 (2021) 10.1073/pnas.2111696118

[35] Moss, D.K., Wilde, A., Lane, J.D.: Dynamic Release of Nuclear RanGTP Triggers TPX2-Dependent Microtubule Assembly During the Apoptotic Execution Phase. J. Cell Sci. 122(5), 644–655 (2009) 10.1242/jcs.037259

[36] Qi, X., Liu, Y., Peng, Y., Fu, Y., Fu, Y., Yin, L., Li, X.: UHRF1 Promotes Spindle Assembly and Chromosome Congression by Catalyzing EG5 Polyubiquitination. J. Cell Biol. 222(11), 202210093 (2023) 10.1083/jcb.202210093

[37] Drechsel, D.N., Hyman, A., Cobb, M.H., Kirschner, M.: Modulation of the Dynamic Instability of Tubulin Assembly by the Microtubule-Associated Protein Tau. Mol. Biol. Cell 3(10), 1141–1154 (1992) 10.1091/mbc.3.10.1141

[38] Dixit, R., Ross, J.L., Goldman, Y.E., Holzbaur, E.L.: Differential Regulation of Dynein and Kinesin Motor Proteins by Tau. Science 319(5866), 1086–1089 (2008) 10.1126/science.1152993

[39] Vershinin, M., Carter, B.C., Razafsky, D.S., King, S.J., Gross, S.P.: Multiple-Motor Based Transport and Its Regulation by Tau. Proc. Natl. Acad. Sci. USA 104(1), 87–92 (2007) 10.1073/pnas.0607919104

[40] Seitz, A., Kojima, H., Oiwa, K., Mandelkow, E.-M., Song, Y.-H., Mandelkow, E.: Single-Molecule Investigation of the Interference Between Kinesin, Tau and MAP2c. EMBO J. (2002) 10.1093/emboj/cdf503

[41] Trinczek, B., Ebneth, A., Mandelkow, E.-M., Mandelkow, E.: Tau Regulates the Attachment/Detachment but Not the Speed of Motors in Microtubule-Dependent Transport of Single Vesicles and Organelles. J. Cell Sci. 112(14), 2355–2367 (1999) 10.1242/jcs.112.14.2355

[42] Ebneth, A., Godemann, R., Stamer, K., Illenberger, S., Trinczek, B., Mandelkow, E.-M., Mandelkow, E.: Overexpression of Tau Protein Inhibits Kinesin-Dependent Trafficking of Vesicles, Mitochondria, and Endoplasmic Reticulum: Implications for Alzheimer’s Disease. J. Cell Biol. 143(3), 777–794 (1998) 10.1083/jcb.143.3.777

[43] Lyu, C., Da Vela, S., Al-Hilaly, Y., Marshall, K.E., Thorogate, R., Svergun, D., Serpell, L.C., Pastore, A., Hanger, D.P.: The Disease-associated Tau35 Fragment Has an Increased Propensity to Aggregate Compared to Full-Length Tau. Front. Mol. Biosci. 8, 779240 (2021) 10.3389/fmolb.2021.779240

[44] Chang, C.-C., Coyle, S.M.: Regulatable Assembly of Synthetic Microtubule Architectures Using Engineered Microtubule-Associated Protein-IDR Condensates. J. Biol. Chem. 300(8) (2024) 10.1016/j.jbc.2024.107544

[45] Maucuer, A., Desforges, B., Joshi, V., Boca, M., Kretov, D.A., Hamon, L., Bouhss, A., Curmi, P.A., Pastré, D.: Microtubules as Platforms for Probing Liquid–Liquid Phase Separation in Cells–Application to RNA-Binding Proteins. J. Cell Sci. 131(11), 214692 (2018) 10.1242/jcs.214692

[46] Dignon, G.L., Zheng, W., Kim, Y.C., Best, R.B., Mittal, J.: Sequence Determinants of Protein Phase Behavior from a Coarse-Grained Model. PLOS Comput. Biol. 14(1), 1005941 (2018) 10.1371/journal.pcbi.1005941

[47] Blas, F.J., MacDowell, L.G., Miguel, E., Jackson, G.: Vapor-Liquid Interfacial Properties of Fully Flexible Lennard-Jones Chains. J. Chem. Phys. 129(14) (2008) 10.1063/1.2989115

[48] Silmore, K.S., Howard, M.P., Panagiotopoulos, A.Z.: Vapour–Liquid Phase Equilibrium and Surface Tension of Fully Flexible Lennard–Jones Chains. Mol. Phys. 115(3), 320–327 (2017) 10.1080/00268976.2016.1262075

[49] Tesei, G., Lindorff-Larsen, K.: Improved Predictions of Phase Behaviour of Intrinsically Disordered Proteins by Tuning the Interaction Range. Open Res. Europ. 2, 94 (2023) 10.12688/openreseurope.14967.2

[50] Uversky, V.N., Gillespie, J.R., Fink, A.L.: Why Are “Natively Unfolded” Proteins Unstructured Under Physiologic Conditions? Proteins: Struct., Funct., Bioinform. 41(3), 415–427 (2000) 10.1002/1097-0134(20001115)41:3⟨415::AID-PROT130⟩3.0.CO;2-7

[51] Guo, C., Alfaro-Aco, R., Zhang, C., Russell, R.W., Petry, S., Polenova, T.: Structural Basis of Protein Condensation on Microtubules Underlying Branching Microtubule Nucleation. Nat. Commun. 14(1), 3682 (2023) 10.1038/s41467-023-39176-z

[52] Ammar Khodja, L., Campanacci, V., Lippens, G., Gigant, B.: The structure of a tau fragment bound to tubulin prompts new hypotheses on tau mechanism and oligomerization. PNAS Nexus 3(11), 487 (2024) 10.1093/pnasnexus/pgae487

[53] Watanabe, S.: Information theoretical analysis of multivariate correlation. IBM Journal of Research and Development 4(1), 66–82 (1960) 10.1147/rd.41.0066

[54] Cohan, M.C., Shinn, M.K., Lalmansingh, J.M., Pappu, R.V.: Uncovering Non-Random Binary Patterns Within Sequences of Intrinsically Disordered Proteins. J. Mol. Biol. 434(2), 167373 (2022) 10.1016/j.jmb.2021.167373

[55] Alfaro-Aco, R., Thawani, A., Petry, S.: Structural Analysis of the Role of TPX2 in Branching Microtubule Nucleation. J. Cell Biol. 216(4), 983–997 (2017) 10.1083/jcb.201607060

[56] Balchand, S.K., Mann, B.J., Titus, J., Ross, J.L., Wadsworth, P.: TPX2 Inhibits Eg5 by Interactions with Both Motor and Microtubule. J. Biol. Chem. 290(28), 17367–17379 (2015) 10.1074/jbc.M114.612903

[57] Kellogg, E.H., Hejab, N.M., Poepsel, S., Downing, K.H., DiMaio, F., Nogales, E.: Near-Atomic Model of Microtubule-Tau Interactions. Science 360(6394), 1242– 1246 (2018) 10.1126/science.aat1780

[58] Tan, R., Lam, A.J., Tan, T., Han, J., Nowakowski, D.W., Vershinin, M., Simó, S., Ori-McKenney, K.M., McKenney, R.J.: Microtubules Gate Tau Condensation to Spatially Regulate Microtubule Functions. Nat. Cell Biol. 21, 1078–1085 (2019) 10.1038/s41556-019-0375-5

[59] Eastman, P., Galvelis, R., Peláez, R.P., Abreu, C.R., Farr, S.E., Gallicchio, E., Gorenko, A., Henry, M.M., Hu, F., Huang, J., et al.: OpenMM 8: Molecular Dynamics Simulation with Machine Learning Potentials. J. Phys. Chem. B 128(1), 109–116 (2023) 10.1021/acs.jpcb.3c06662

[60] Izaguirre, J.A., Sweet, C.R., Pande, V.S.: Multiscale Dynamics of Macromolecules Using Normal Mode Langevin. Pac. Symp. Biocomput. 15, 240–251 (2010) 10.1142/97898142952910026

[61] Chow, K.-H., Ferguson, D.M.: Isothermal-Isobaric Molecular Dynamics Simulations with Monte Carlo Volume Sampling. Comput. Phys. Commun. 91(1-3), 283–289 (1995) 10.1016/0010-4655(95)00059-O

[62] Åqvist, J., Wennerstrüom, P., Nervall, M., Bjelic, S., Brandsdal, B.O.: Molecular Dynamics Simulations of Water and Biomolecules with a Monte Carlo Constant Pressure Algorithm. Chem. Phys. Lett. 384(4-6), 288–294 (2004) 10.1016/j.cplett.2003.12.039

