## Supporting Information for "Molecular Grammar of Microtubule-Wetting Condensates"

#### S1 Inherently disordered proteins (IDPs) studied

This section briefly discusses the choice and relevance of the IDPs studied in this work, and how their structures were obtained. This study aims to investigate the capability of coarse-grained molecular simulations to predict the selective wetting of IDPs to wet MTs. Therefore, it was prudent to select IDRs that are reported to condense on MTs. To include negative controls in the study, we also studied IDRs that do not wet the surface of MTs according to experimental investigation reports. In total, eight IDP sequences were identified from the literature (see table S2). The rest of this section describes these sequences and how their initial configurations were generated.

The N-terminal domain (NTD) of the LEM2 (LEM2-NTD) has been reported to condense on MTs, enabling them to carry out functions during cell division [S1]. The current study focuses only on inherently disordered regions (IDRs, or low complexity domains (LCDs)). Therefore, the ordered domains of LEM2-NTD were excluded, and the disordered domain from residues 71-208 was sliced from the AlphaFold structure Q8NC56 [S2, S3] for simulating the LCD region of LEM2 (LEM2-LCD).

The whole sequence of Tau 2N4R was obtained from UniProt entry P10636-8.Tau35 is a 35kDa fragment (from E187 to L441) in the C-terminal region of Tau 2N4R [S4], and is reported to have a higher propensity to aggregate when compared to the full-length Tau, and has all the MT-binding repeats [S5]. The necessary domain (E187-L441) containing 255 residues was sliced to generate the initial configuration for simulations. Necessary mutations were applied to the sliced AlphaFold [S2, S3] prediction (AF-P10636-F1-v4) of the Tau protein using CHARMM-GUI [S6, S7] to generate the initial all-atom structure for the simulations of Tau35.

TPX2 (targeting protein for Xklp2) is a protein that stimulates branching MT nucleation. They play an important role during apoptosis, in the normal assembly of microtubules. They are also required for chromatin and/or kinetochore-dependent nucleation of microtubule [S8, S9, S10]. The region in the C-terminal part of TPX2 (Xenopus laevis) protein (residues 477–716) constitutes the minimal region of TPX2 which is capable of stimulating branching microtubule nucleation [S11, S12]. For the simulations in the current work, the initial all-atom configuration for human TPX2 (residues 477-716) was obtained from the AlphaFold database (AF-Q9ULW0-F1-v4) of human TPX2 and was treated as an IDP in simulations. This domain of TPX2 is referred to as TPX2 throughout the main article and the supporting information file.

The LCD of FUS is reported to exhibit LLPS [S13] but was observed not to bind to the surface of MTs [S14]. The initial configuration was obtained from the AlphaFold database (AF-P35637-F1-v4) [S2, S3]. The LCD domain of the protein (FUS-LCD) used in simulations by Farag et al. [S13] was created by slicing residues 1-214.

Chang et al [S14] also reported that DDX4 and LAF1 do not bind to the surface of MTs. The region from residue 1-236 in the NTD of DDX4 (DDX4-NTD) was used in their study and was observed to not wet the surface of MTs in experiments. For the current work, we used the same segment of the DDX4 protein. The initial all-atom 3D structure for simulations was obtained from the Alphafold database (AF-Q9NQI0-F1-v4) [S2, S3], and the domain of interest was sliced from the protein.

As mentioned earlier, Chang et al. also reported the non-MT-binding nature of LAF1-NTD. They worked with the domain of LAF1 from residue 1-200. Another work that studied the liquid-liquid phase separation of the domain of LAF1 rich in the Arginine-Glycine-Glycine motif was reported in the literature by Schuster et al. [S15]. This domain of the LAF1, the LAF1-RGG (residues 1-176) was used in our simulations. The initial all-atom structure was obtained from AlphaFold database [S2, S3] (AF-Q9M0K4-F1-v4). The RGG-domain (1-176) was sliced to generate the initial all-atom structure of LAF1-RGG.

#### S1.1 Sequences

The amino acid sequences of all the IDRs described above are listed in this section. The residue numbers used to identify specific residues in the sequences in this work are based on 0-based indexing of the sequence.

##### LEM2-LCD

```
DAPLRARPAA ASPRAEPWLS QPASGSAYAT PGAYGDIRPS AASWVGSRGL AYPARPAQLR RRASVRGSSE
EDEDARTPDR ATQGPGLAAR RWWAASPAPA RLPSSLLGPD PRPGLRATRA GPAGAARARP EVGRRLER
```

##### Tau 2N4R

```
MAEPRQEFEV MEDHAGTYGL GDRKDQGGYT MHQDQEGDTD AGLKESPLQT PTEDGSEEPG SETSDAKSTP
TAEDVTAPLV DEGAPGKQAA AQPHTEIPEG TTAEEAGIGD TPSLEDEAAG HVTQARMVSK SKDGTGSDDK
KAKGADGKTK IATPRGAAPP GQKGQANATR IPAKTPPAPK TPPSSGEPPK SGDRSGYSSP GSPGTPGSRS
RTPSLPTPPT REPKKVAVVR TPPKSPSSAK SRLQTAPVPM PDLKNVKSKI GSTENLKHQP GGGKVQIINK
KLDLSNVQSK CGSKDNIKHV PGGGSVQIVY KPVDLSKVTS KCGSLGNIHH KPGGGQVEVK SEKLDFKDRV
QSKIGSLDNI THVPGGGNKK IETHKLTFRE NAKAKTDHGA EIVYKSPVVS GDTSPRHLSN VSSTGSIDMV
DSPQLATLAD EVSASLAKQG L
```

##### Tau35

```
EPPKSGDRSG YSSPGSPGTP GSRSRTPSLP TPPTREPKKV AVVRTPPKSP SSAKSRLQTA PVPMPDLKNV
KSKIGSTENL KHQPGGGKVQ IINKKLDLSN VQSKCGSKDN IKHVPGGGSV QIVYKPVDLS KVTSKCGSLG
NIHHKPGGGQ VEVKSEKLDF KDRVQSKIGS LDNITHVPGG GNKKIETHKL TFRENAKAKT DHGAEIVYKS
PVVSGDTSPR HLSNVSSTGS IDMVDSPQLA TLADEVSASL AKQGL
```

##### FUS-Tau35

```
MASNDYTQQA TQSYGAYPTQ PGQGYSQQSS QPYGQQSYSG YSQSTDTSGY GQSSYSSYGQ SQNTGYGTQS
TPQGYGSTGG YGSSQSSQSS YGQQSSYPGY GQQPAPSSTS GSYGSSSQSS SYGQPQSGSY SQQPSYGGQQ
QSYGQQQSYN PPQGYGQQNQ YNSSSGGGGG GGGGGNYGQD QSSMSSGGGS GGGYGNQDQS GGGGSGGYGQ
QDRGEPPKSG DRSGYSSPGS PGTPGSRSRT PSLPTPPTRE PKKVAVVRTP PKSPSSAKSR LQTAPVPMPD
LKNVKSKIGS TENLKHQPGG GKVQIINKKL DLSNVQSKCG SKDNIKHVPG GGSVQIVYKP VDLSKVTSKC
GSLGNIHHKP GGGQVEVKSE KLDFKDRVQS KIGSLDNITH VPGGGNKKIE THKLTFRENA KAKTDHGAEI
VYKSPVVSGD TSPRHLSNVS STGSIDMVDS PQLATLADEV SASLAKQGL
```

##### TPX2

```
KVLPITVPKS PAFALKNRIR MPTKEDEEED EPVVIKAQPV PHYGVPFKPQ IPEARTVEIC PFSFDSRDKE
RQLQKEKKIK ELQKGEVPKF KALPLPHFDT INLPEKKVKN VTQIEPFCLE TDRRGALKAQ TWKHQLEEEL
RQQKEAACFK ARPNTVISQE PFVPKKEKKS VAEGLSGSLV QEPFQLATEK RAKERQELEK RMAEVEAQKA
QQLEEARLQE EEQKKEELAR LRRELVHKAN
```

##### FUS-LCD

```
MASNDYTQQA TQSYGAYPTQ PGQGYSQQSS QPYGQQSYSG YSQSTDTSGY GQSSYSSYGQ SQNTGYGTQS
TPQGYGSTGG YGSSQSSQSS YGQQSSYPGY GQQPAPSSTS GSYGSSSQSS SYGQPQSGSY SQQPSYGGQQ
QSYGQQQSYN PPQGYGQQNQ YNSSSGGGGG GGGGGNYGQD QSSMSSGGGS GGGYGNQDQS GGGGSGGYGQ
QDRG
```

##### LAF1-RGG

```
MESNQSNNGG SGNAALNRGG RYVPPHLRGG DGGAAAAASA GGDDRRGGAG GGGYRRGGGN SGGGGGGGYD
RGYNDNRDDR DNRGGSGGYG RDRNYEDRGY NGGGGGGGNR GYNNNRGGGG GGYNRQDRGD GGSSNFSRGG
YNNRDEGSDN RGSGRSYNND RRDNGGDGLE HHHHHH
```

**DDX4-NTD**

```
MGDEDWEAEI NPHMSSYVPI FEKDRYSGEN GDNFNRTPAS SSEMDDGPSR RDHFMKSGFA SGRNFGNRDA
GECNKRDNTS TMGGFGVGKS FGNRGFSNSR FEDGDSSGFW RESSNDCEDN PTRNRGFSKR GGYRDGNNSE
ASGPYRRGGR GSFRGCRGGF GLGSPNNDLD PDECMQRTGG LFGSRRPVLS GTGNGDTSQS RSGSGSERGG
YKGLNEEVIT GSGKNSWKSE AEGGES
```

#### S1.2 Sequence Properties

Sequence descriptors have been reported in the literature to quantify the features of amino-acid sequences, which are often important in the physical behavior of the IDRs (Table S1).

The kappa ($\kappa$) paremeter was developed by Pappu and coworkers [S16] to quantify the sequence patterning of oppositely charged residues. The patterning of charged residues can directly contribute to global compaction or expansion of IDPs [S17]. The value of $\kappa$ lies between 0 and 1. For a fixed composition of aminoacids, a lower value of $\kappa$ indicates that the oppositely charged residues are well mixed within the sequence. It has also been reported that the fraction of charged residues (FCR) and the patterning of charged residues affect the size, shape, and conformational fluctuations of certain IDRs [S16]. The mean Uversky-hydrophobicity of the chain and the mean net charge has been argued to delineate putative IDPs and autonomously foldable proteins [S18]. Omega ($\omega$) quantifies the linear segregation (or mixing) of prolines and charged residues versus all other residues [S19], and is relevant for IDRs that are rich in charged residues and proline [S19]. We also calculated the binary mixing between different residue types, and is discussed in section S4.3 in the document. The diversity of IDRs in terms of their properties is demonstrated using the parameters listed in table S1.

| Sequence Property | LEM2-LCD | Tau 2N4R | TPX2 | Tau 35 | FUS-Tau35 | FUS-LCD | DDX4-NTD | LAF1-RGG |
|---|---|---|---|---|---|---|---|---|
| Net charge | +11 | +2 | +6 | +17 | 13 | -4 | -4 | +3 |
| Number of residues | 138 | 441 | 240 | 255 | 469 | 214 | 236 | 176 |
| Net charge per residue (NCPR) | +0.07971 | +0.004535 | +0.025 | +0.066667 | +0.0277186 | -0.018692 | -0.016949 | 0.017045 |
| f+ (fraction of positive charged residues) | 0.166667 | 0.131519 | 0.2 | 0.152941 | 0.0852878 | 0.004673 | 0.135593 | 0.136364 |
| f- (fraction of negatively charged residues) | 0.086957 | 0.126984 | 0.175 | 0.086275 | 0.0575693 | 0.023364 | 0.152542 | 0.119318 |
| Fraction of charged residues (FCR) | 0.253623 | 0.258503 | 0.375 | 0.239216 | 0.142857 | 0.028037 | 0.288136 | 0.255682 |
| kappa | 0.220243 | 0.184127 | 0.15002 | 0.126845 | 0.135583 | 0.207515 | 0.237054 | 0.101418 |
| Omega | 0.119305 | 0.114456 | 0.097852 | 0.129724 | 0.397879 | 1.133621 | 0.13291 | 0.317948 |
| Uversky hydropathy | 0.416908 | 0.403578 | 0.392824 | 0.420741 | 0.384009 | 0.340239 | 0.358239 | 0.307071 |

Table S1: Sequence properties of the IDRs studied in this work.

### S2 Homogeneous Liquid-Liquid Phase Separation

| IDR | Residues | Number of Chains | Box size ($\mathbf{L}_x \times \mathbf{L}_y \times \mathbf{L}_z$) nm |
|---|---|---|---|
| Tau 2N4R | 441 | 100 | 25.0 × 25.0 × 120.7 |
| Tau35 | 255 | 100 | 17.0 × 17.0 × 261.1 |
| LEM2-LCD | 138 | 100 | 15.0 × 15.0 × 335.4 |
| TPX2 | 240 | 100 | 17.0 × 17.0 × 261.1 |
| FUS-Tau35 | 469 | 100 | 25.0 × 25.0 × 120.7 |
| FUS-LCD | 214 | 100 | 17.0 × 17.0 × 261.1 |
| LAF1-RGG | 176 | 100 | 15.0 × 15.0 × 335.4 |
| DDX4-NTD | 236 | 100 | 17.0 × 17.0 × 261.1 |

Table S2: IDPs used in the study. LEM2-LCD, Tau 2N4R, and TPX2 have been reported to condense on MT [S1, S5, S20, S21]. Whereas, FUS-LCD, LAF1-RGG and DDX4 have been reported not to have an affinity towards MT [S14]. We also included Tau35 fragment, and a combined FUS-Tau35 IDR in the study.

'Slab simulations' have been reported in the literature [S22, S23, S24, S25] to study and report phase separation of IDRs in simulations. The IDRs are initially brought to the center of the box by applying a harmonic spring to each bead. The system is then allowed to evolve freely at a specified temperature (Fig. S1a provides a visualization of the slab simulation of LEM2-LCD). In the simulation protocol, if there is a phase separation, the high-density phase with surfaces along the x-y plane is simulated in equilibrium with the low-density phase. The density profile along normal to the phase surfaces (z-axis) was estimated as shown in Fig. S1b. Before the density profile is estimated, the trajectory is centered on the 'slab' in each frame. The definition of the slab as the "cluster with the largest number of chains" used by Dignon et al [S22] was used. Clustering was done based on a center-of-mass distance criterion with a distance cut-off of 5 nm.

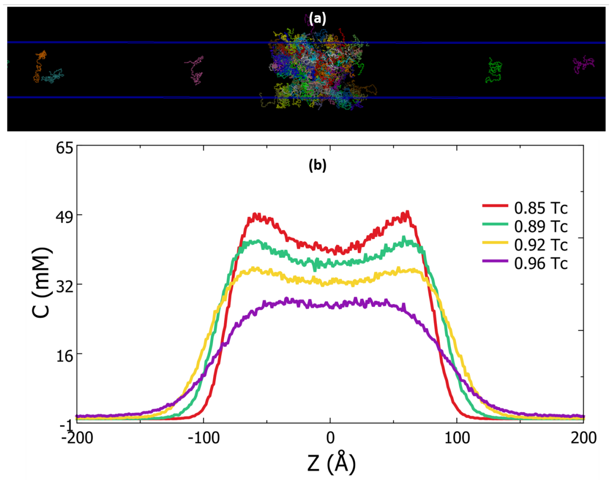


Figure S1: (a) A visualisation of the slab simulation of LEM2-LCD IDR. For density analysis, the slab is maintained at the center of the box in the trajectory by identifying the largest cluster and translating it to the center of the box. (b) The density profile was measured along the length of the box to determine the concentrations of the high-density and low-density phases.

The critical temperature $T_c$ was obtained by fitting

$$\rho_h - \rho_l = A\,(T_c - T)^{\beta} \tag{S1}$$

where $\rho_h$ and $\rho_l$ represent the density of the high and the low-density phases (respectively), $T_c$ is the critical temperature, A is the protein specific fitting parameter, and $\beta$ is set to 0.325 (universality class of 3D Ising model [S26]). The lowest temperature for fitting was the lowest temperature where $\rho_l$ is non-zero. The highest temperature for fitting was determined by minimizing the error while fitting equation S1. Temperatures greater than $T_c$ were not considered. The values of $\Delta\rho$ ($\rho_h$ - $\rho_l$) against temperature are shown in Fig. S2 for all the IDRs studied in this work.

#### S2.1 Residue-Residue contact distances

The residue-residue contact distances were determined from the total energy of non-bonded interactions between the two residues as defined by the CALVADOS3 potential energy function [S27]. If the total interaction energy was less than -0.1 kT, two residues were identified to be in contact. The total potential energy between two residues is given by equation S2

$$U_{\text{Total}} = U_{\text{AH}} + U_{\text{DH}} \tag{S2}$$

where, $U_{AH}$ represents the truncated and shifted Ashbaugh-Hatch potential (to model van der Waals interactions) and $U_{DH}$ represents the Debye-Hückel potential (to model salt-screened electrostatic interactions). The Ashbaugh-Hatch potential is represented in equation S3

$$U_{\text{AH}}(r) = \begin{cases} U_{\text{LJ}}(r) - \lambda U_{\text{LJ}}(r_c) + \epsilon(1-\lambda), & r \le 2^{1/6}\sigma \\ \lambda\left[U_{\text{LJ}}(r) - U_{\text{LJ}}(r_c)\right], & 2^{1/6}\sigma < r \le r_c \\ 0, & r > r_c \end{cases} \tag{S3}$$

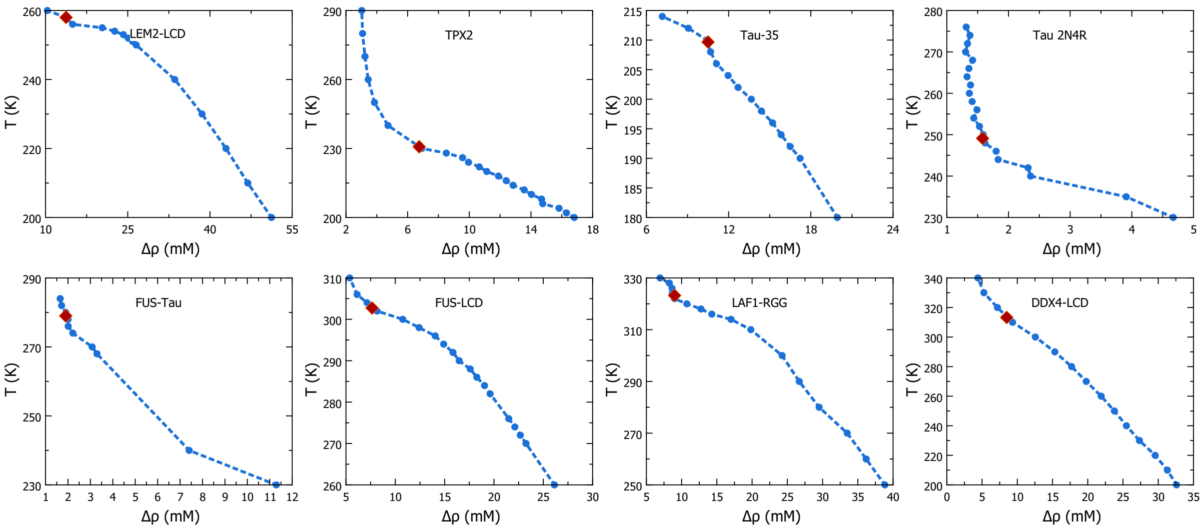


Figure S2: Phase data of all IDR used in the study. The difference between the densities if the high density and low-density phases is plotted against the temperature. The critical temperature of the system, estimated using eqn S1 is shown in red color.

where, $\lambda$ is the 'stickiness' parameter, $r_c$ is the truncation distance (set to 2.2 nm), and $\epsilon$ = 0.8368 kJ $mol^{-1}$. $U_{LJ}$ describes the Lennard-Jones potential, shown in equation S4.

$$U_{\mathrm{LJ}}(r) = 4\epsilon\left[\left(\frac{\sigma}{r}\right)^{12} - \left(\frac{\sigma}{r}\right)^{6}\right] \tag{S4}$$

$\sigma$ in equations S3 and S4 represents the bead size of residues, and both $\sigma$ and $\lambda$ are the arithmetic averages of residue-specific bead sizes and 'stickiness' values, respectively.
The Debye-Hückel potential used to describe electrostatic interactions in the system is shown in equation S5

$$U_{\mathrm{DH}}(r) = \frac{q_i q_j e^2}{4\pi\epsilon_0\epsilon_r}\left[\frac{\exp(-r/D)}{r}\right] \tag{S5}$$

where q is the residue's charge, and e is the charge of an electron. The Debye length D is estimated as $\sqrt{1/(8\pi B c_s)}$, where $C_s$ is the ionic strength, and $B(\epsilon_r)$ is the Bjerrum length. Temperature dependant relative permittivity ($\epsilon_r$) is estimated as

$$\epsilon_r(\mathrm{T}) = \frac{5321}{\mathrm{T}} + 233.76 - 0.9297\times\mathrm{T} + 1.417\times10^{-3}\times\mathrm{T}^2 - 8.292\times10^{-7}\times\mathrm{T}^3 \tag{S6}$$

For each set of residue pairs, $r_{min}$ and $r_{max}$ are estimated based on the interaction energy as described. Two residues are defined to be in contact if the distance between them (r) is such that $r_{min} < r < r_{max}$.

### S3 Microtubule-Wetting

#### S3.1 Structure of the Microtubule

The all-atom structure of the microtubule assembly was obtained from protein data bank ID 3J2U [S28]. The missing coordinates of residues 35-60 of $\alpha$-tubulin were generated using CHARMM GUI's PDB Reader & Manipulator [S29]. The structure in the PDB file requires an additional 3 $\alpha$- and $\beta$- tubulin pairs to complete the periodic cylindrical structure along the axis of the MT-assembly (see Fig. S3). These were inserted by translating the $\alpha$- and $\beta$- tubulins vertically above them. For identifying the surface atoms of the Microtubule, the solvent accessible surface area (SASA) was used. In the coarse-grained structure, if the SASA was greater than 55 Å$^2$, the residue was considered to be on the surface. A cut-off 0.5 nm was used to identify the residues in the immediate vicinity of the residue. Only the residues that had surface neighbors within this distance were used to characterize the neighborhood composition. ALL the residue indices reported in the analysis part of this work are based on a 0-based indexing system.

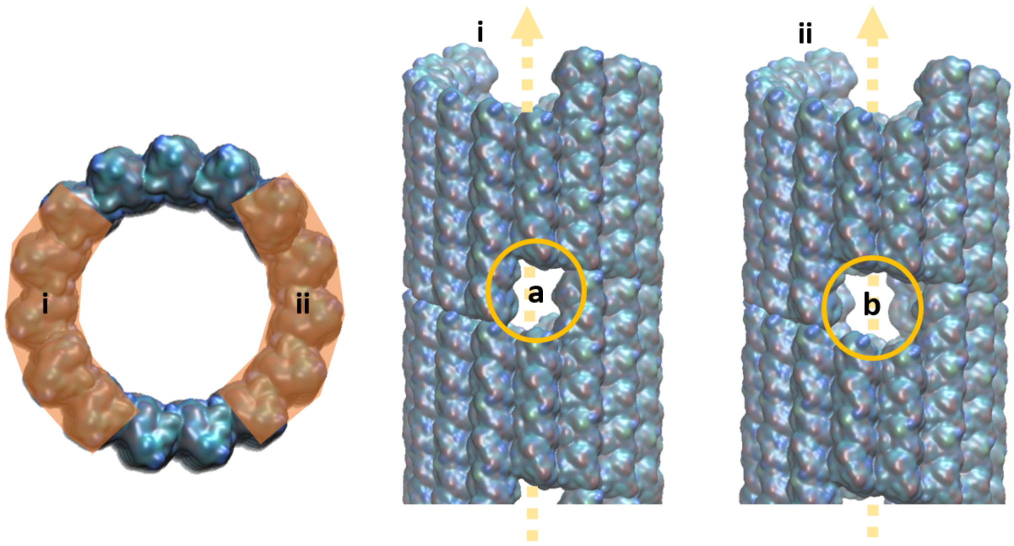


Figure S3: Missing pairs of $\alpha$- and $\beta$- tubulins for completing the periodic cylindrical structure of the microtubule assembly reported in PDB ID 3J2U. Missing pairs of tubulins on two diametrically opposite surfaces of the MT assembly (surfaces (i) and (ii) shown in the figure) create 'holes' on the periodic MT surface. The holes shown as (a) and (b) in the present figure were formed by the absence of one and two $\alpha - \beta$ tubulin pairs, respectively. The periodic structure was restored by adding the coordinates of the required number of tubulin pairs (for each hole). The cordinates of the newly added tubulin molecules were determined by shifting the positions of pairs of $\alpha-$ and $\beta$- tubulins above the 'holes' along the axis (yellow dashed line) of the microtubule by a distance equivalent to the spacing between two $\alpha$- (or $\beta$) tubulins (along the axis of the MT). The cylindrical structure thus generated had 45 pairs of $\alpha$ and $\beta$ tubulins and was used for simulations in the study.

#### S3.2 Wall Potential

The work aims to understand the wetting activity of IDRs on the outer surface of MT. Therefore, it is important to perform simulations where the concentration of the IDR outside the MT surface is maintained constant. In unbiased simulations of IDRs (especially ones that wet the surface of the MT), it was observed that IDRs can 'leak' into the inner hollow region of the MT, altering their concentration outside the MT during the simulation (See Fig. S4a). Maintaining the IDR concentration exterior to the MT could be achieved through a biasing potential that satisfies the following criteria: (i) The potential should be experienced only by particles of the IDR chains, and not the atoms of the MT, (ii) the potential should not affect particles that are at a radial distance r $\geq$ the inner surface radius of the MT. (iii) The potential should provide an energy penalty for particles that are moving inside the inner-MT surface (see Fig. S4b). To achieve these objectives, a biasing wall potential represented by (equation S7) was applied at the inner surface of the micro-tubule.

$$\mathrm{U_{wall}}(r_{xy}) = \begin{cases} \frac{1}{2}k(r_c - r_{xy})^2, & \text{if } r_{xy} < r_c \\ 0, & \text{if } r_{xy} \geq r_c \end{cases} \tag{S7}$$

where $r_{xy} = \sqrt{(x - x_c)^2 + (y - y_c)^2}$ is the radial distance from the center of the MT in the $xy$-plane. $k$ is the stiffness of the harmonic wall potential. $r_c$ is the cutoff radius beyond which the wall exerts no influence.$r_c$ was set to the inner surface radius of the MT ($\sim$ 9 nm). When the distance $r_{xy}$ is less than the cutoff $r_c$, the particle experiences a harmonic repulsive force. The wall has no effect for $r_{xy} \geq r_c$.

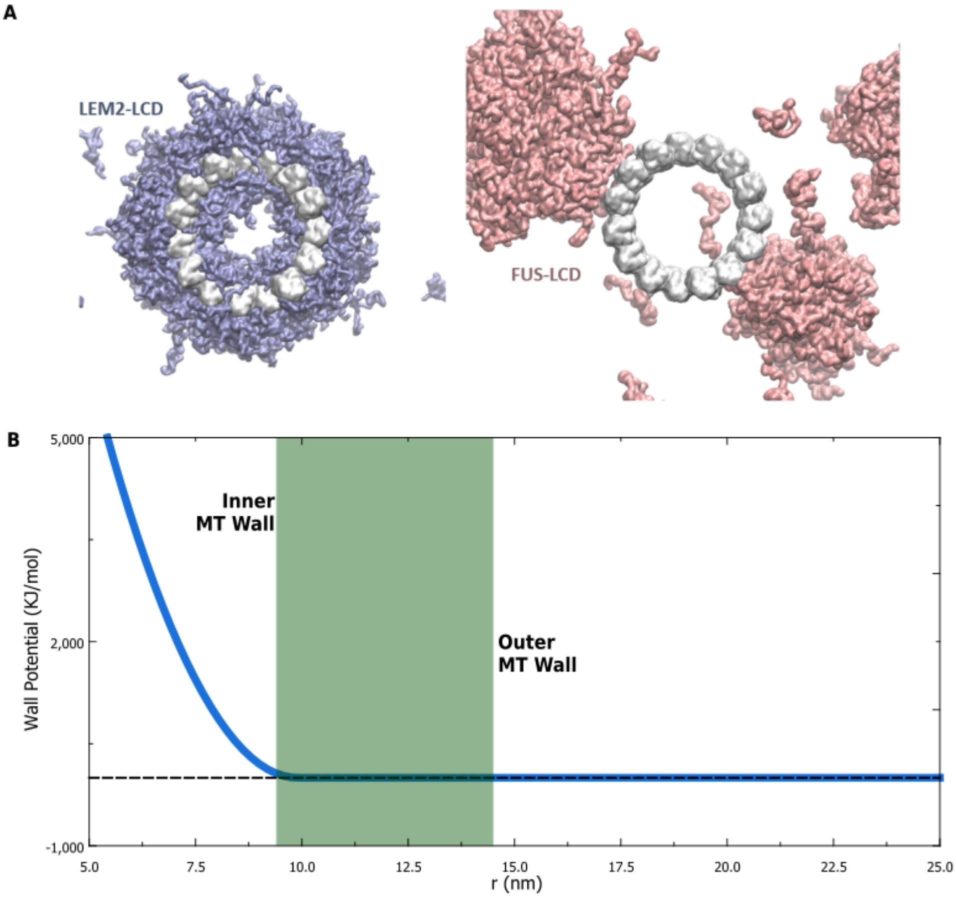


Figure S4: (a) MT-IDR condensate simulations for LEM2-LCD (which wets the MT surface) and FUS-LCD (which does not wet the MT surface) when no external biasing potentials were applied to the system. The IDRs that wet the MT surface were observed to 'leak' into the inner region of MT (as demonstrated with LEM2-LCD in the figure), thereby affecting the concentration of IDRs on the surface, and in the region exterior to the MT (b) Profile of the biasing wall potential applied at the inner surface of the MT. The wall potential provides a harmonic repulsion to IDR atoms when they are at the inner surface of the MT. It does not affect any other particle in the system. This biasing potential prevents the 'leakage' of IDRs into the MT and helps to maintain the concentration of the IDRs outside the MT at specified levels. The wall-potential drops to zero at the inner surface of MT, and therefore does not alter/bias the interaction of IDR with the outer surface of the MT.

##### S3.3 Simulation Protocol for MT-IDR Simulations

We employed 'escape simulations' to study the selectivity in the wetting of MT surface by IDRs. The number of IDR proteins inserted in the system was determined so that the system can have 20% of volume (external to the MT surface) with the high-density condensate of the IDR, and 80% of the volume with IDR in low-density phase. The thickness of the condensate on was assumed to be 10 nm on the surface of MT (for determining the number of proteins in the system).

The residues of the IDR chains were pulled to the surface of the MT with a harmonic spring. The biasing force was applied for 100ns, in an NVT ensemble. The system temperature was maintained using the Langevin integrator [S30] with a friction coefficient of 0.01/picoseconds. Subsequently, the biasing force was removed, and the IDR molecules were allowed to move without other restraints. A wall potential was applied on the inner surface of the MT as discussed in Section S3.2 to prevent the leakage of the IDRs inside the MT cylinder. Position restraints were applied to the residues of the MT atoms to maintain its structure. The production simulations were run for 10 $\mu$S to test the wetting of the MT surface. Fig. S5 illustrates the selectivity in MT-surface wetting by IDR condensates.

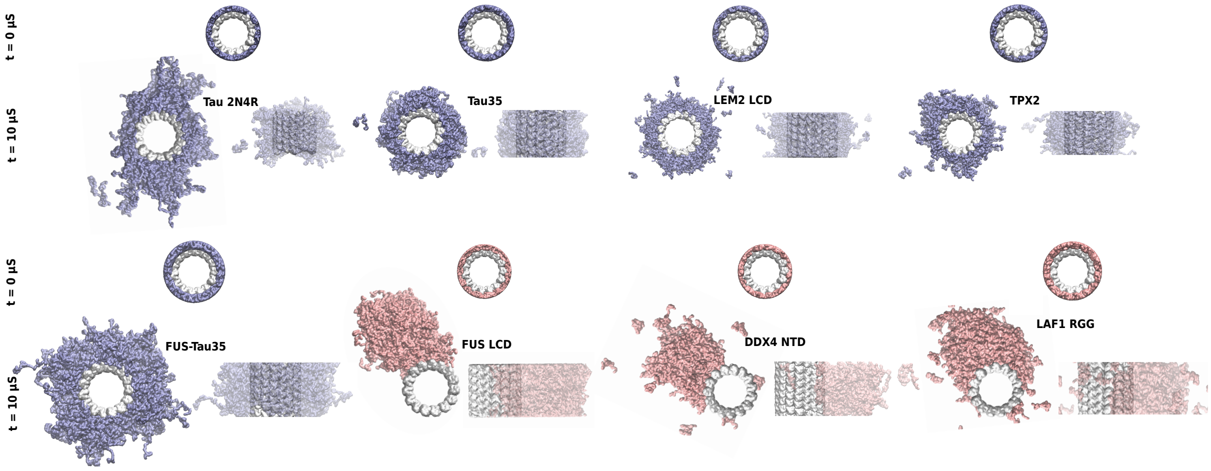


Figure S5: The visualisation of IDR condensates on the surface of MT simulations. Tau 2N4R, Tau35, LEM2 LCD, TPX2, and the combined FUS-Tau35 protein are observed to wet the surface of the MT. On the other hand, FUS LCD, DDX4 NTD, and LAF1 RGG condensates do not wet the surface of the MT.

#### S4 Interactions

##### S4.1 IDR-IDR Interactions

The interactions within the bulk of the condensates were studied using NPT simulations IDR chains at 1 bar. The residue-wise contact maps of all IDRs are shown in Figures S6-S13. Later in Fig. S15, we also show the 'active' blocks in the IDR chains responsible for the stabilisation of the IDR condensate.

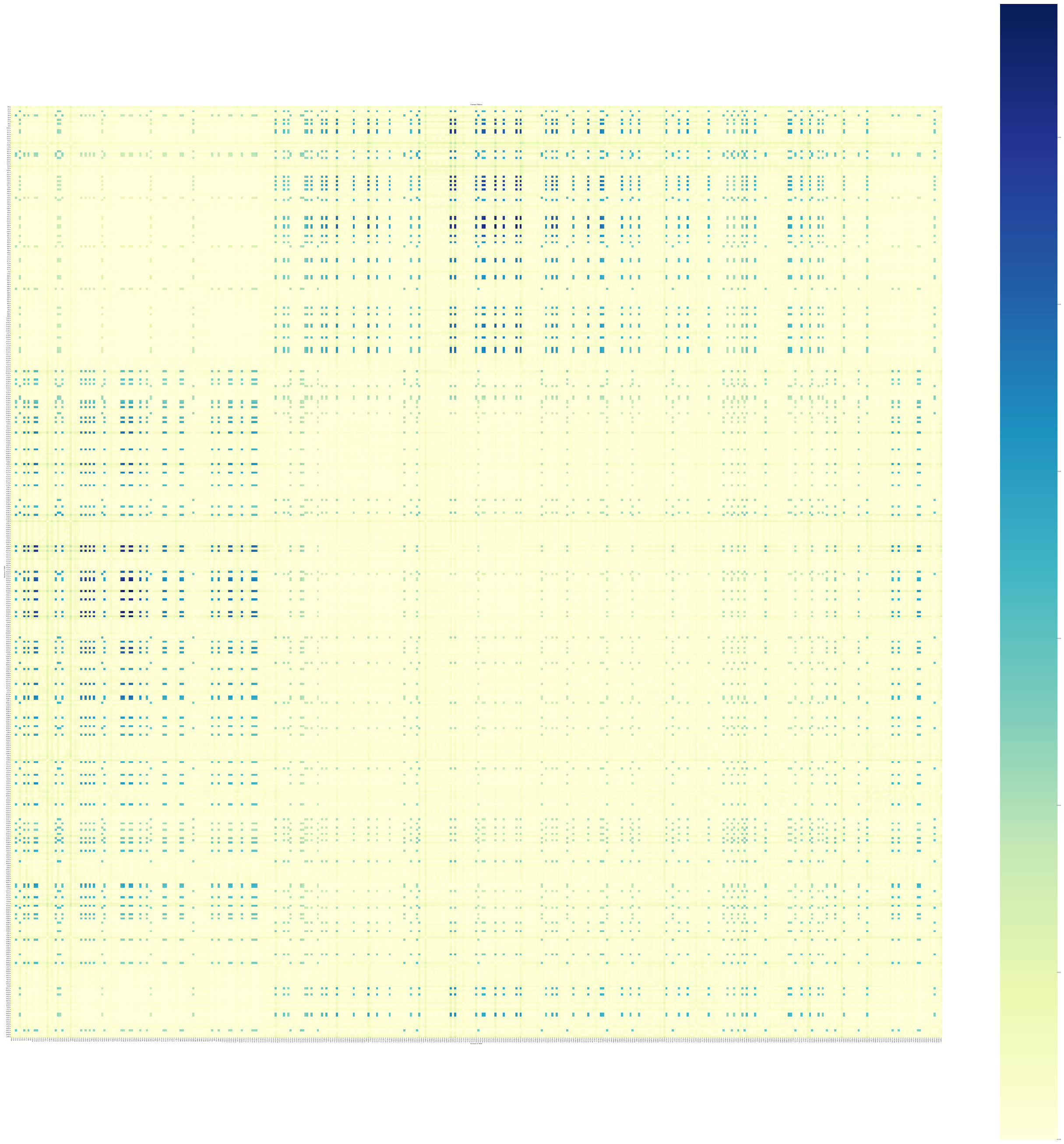

Figure S6: The probability of contact formation between residues of the Tau 2N4R IDR in the bulk of the condensate.

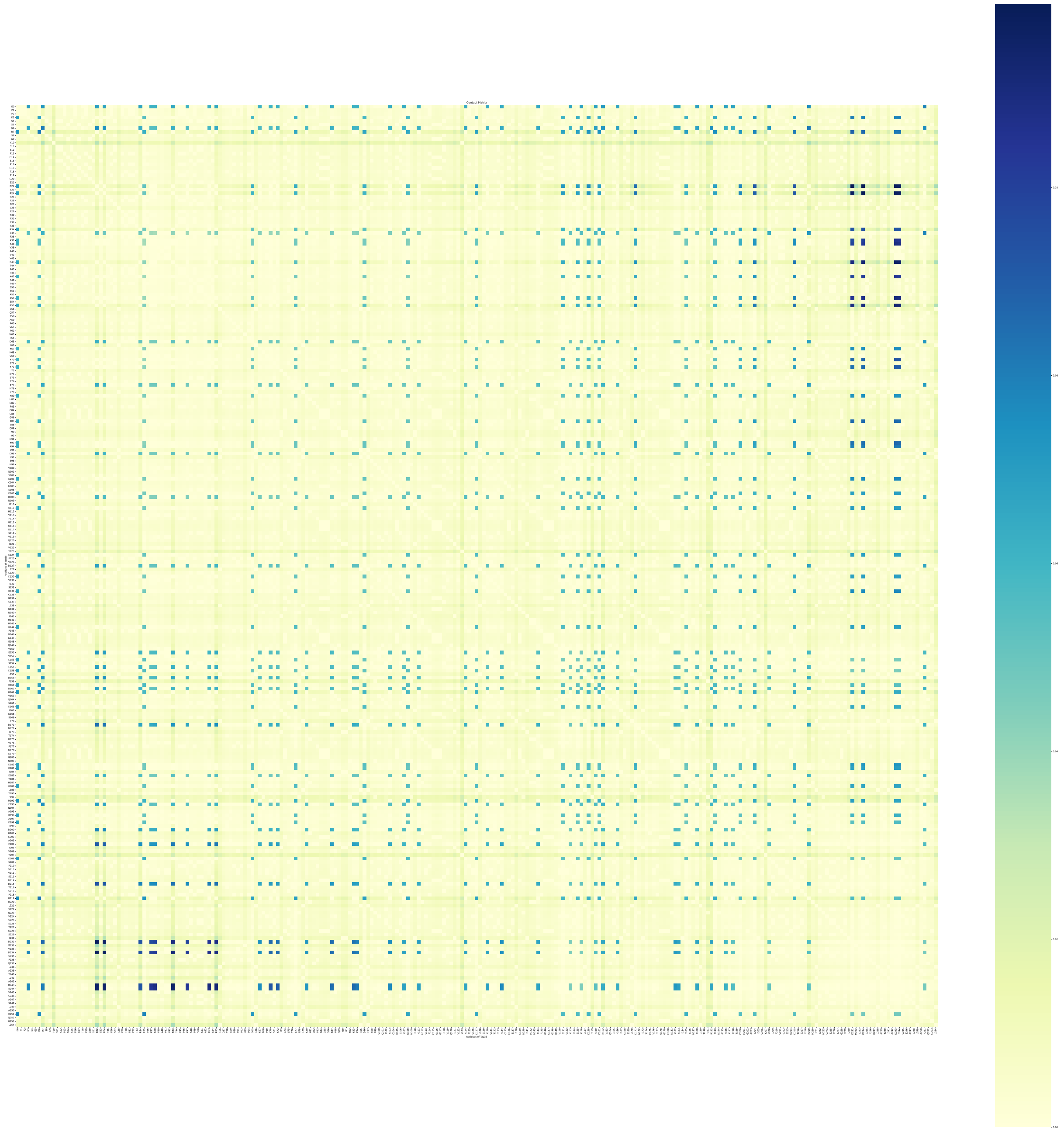

Figure S7: The probability of contact formation between residues of the Tau35 IDR in the bulk of its condensate.

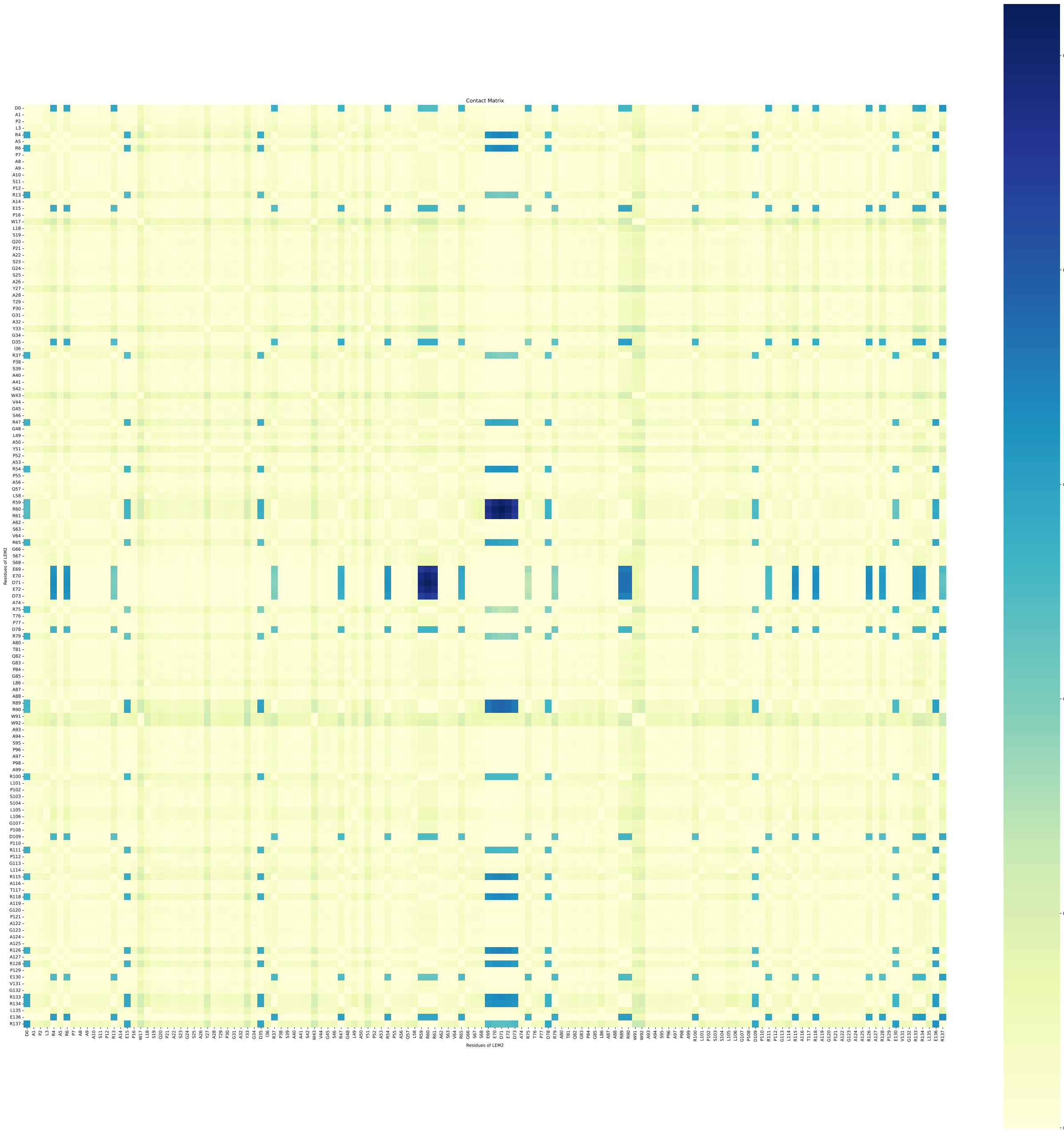


Figure S8: The probability of contact formation between residues of the LEM2 LCD in the bulk of its condensate.

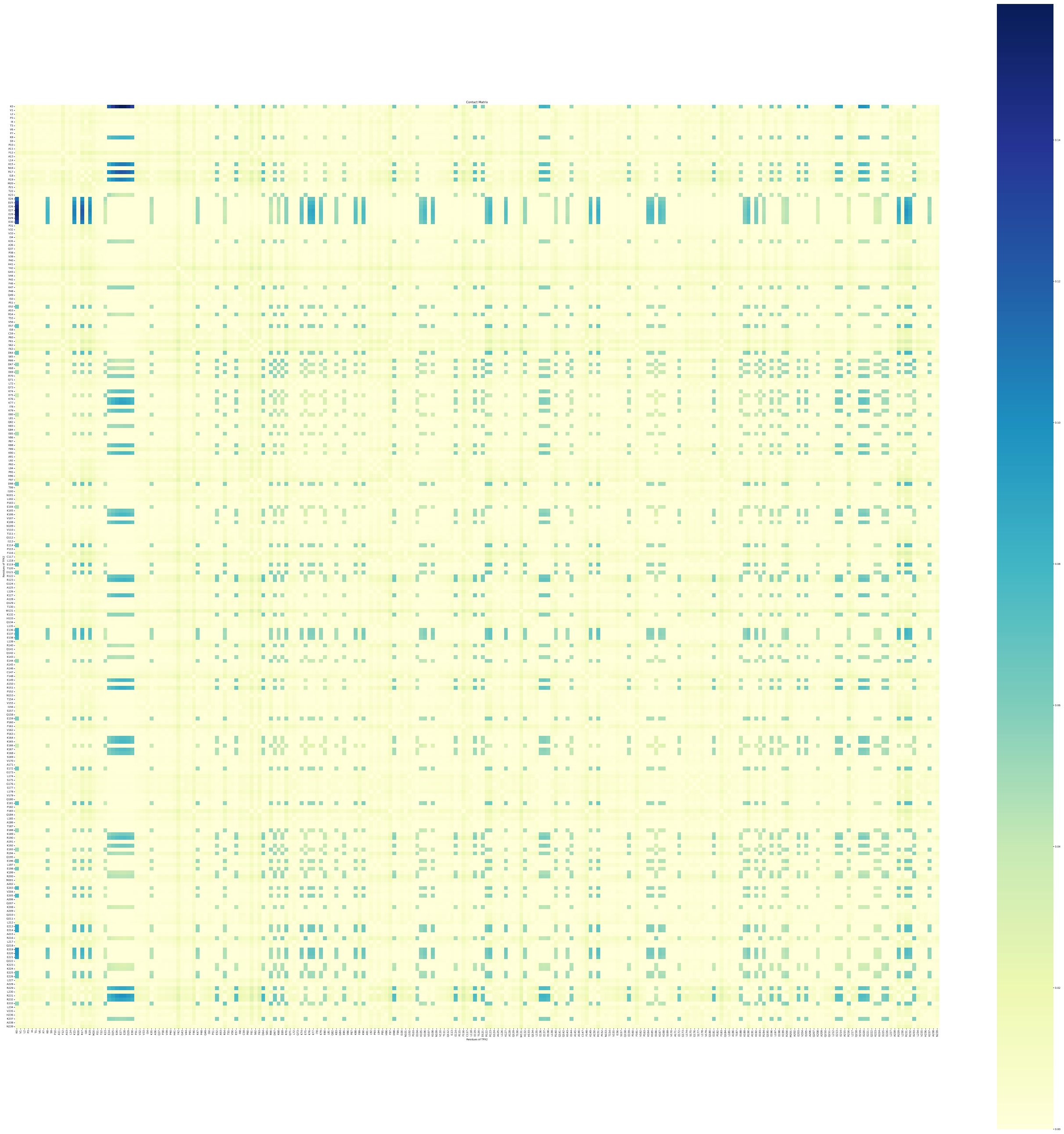

Figure S9: The probability of contact formation between residues of the TPX2 IDR in the bulk of its condensate.

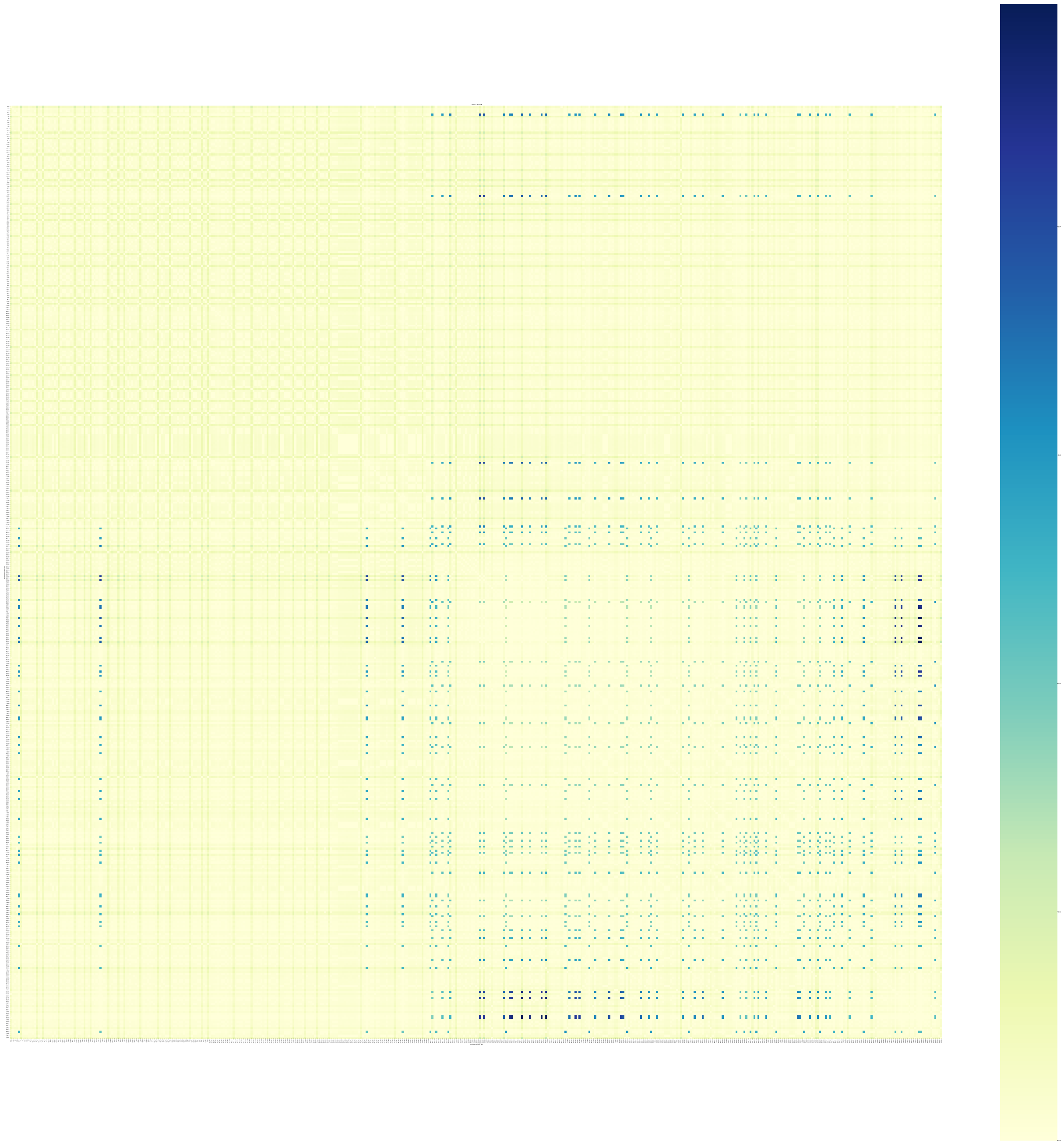

Figure S10: The probability of contact formation between residues of the FUS-Tau35 IDR in the bulk of its condensate.

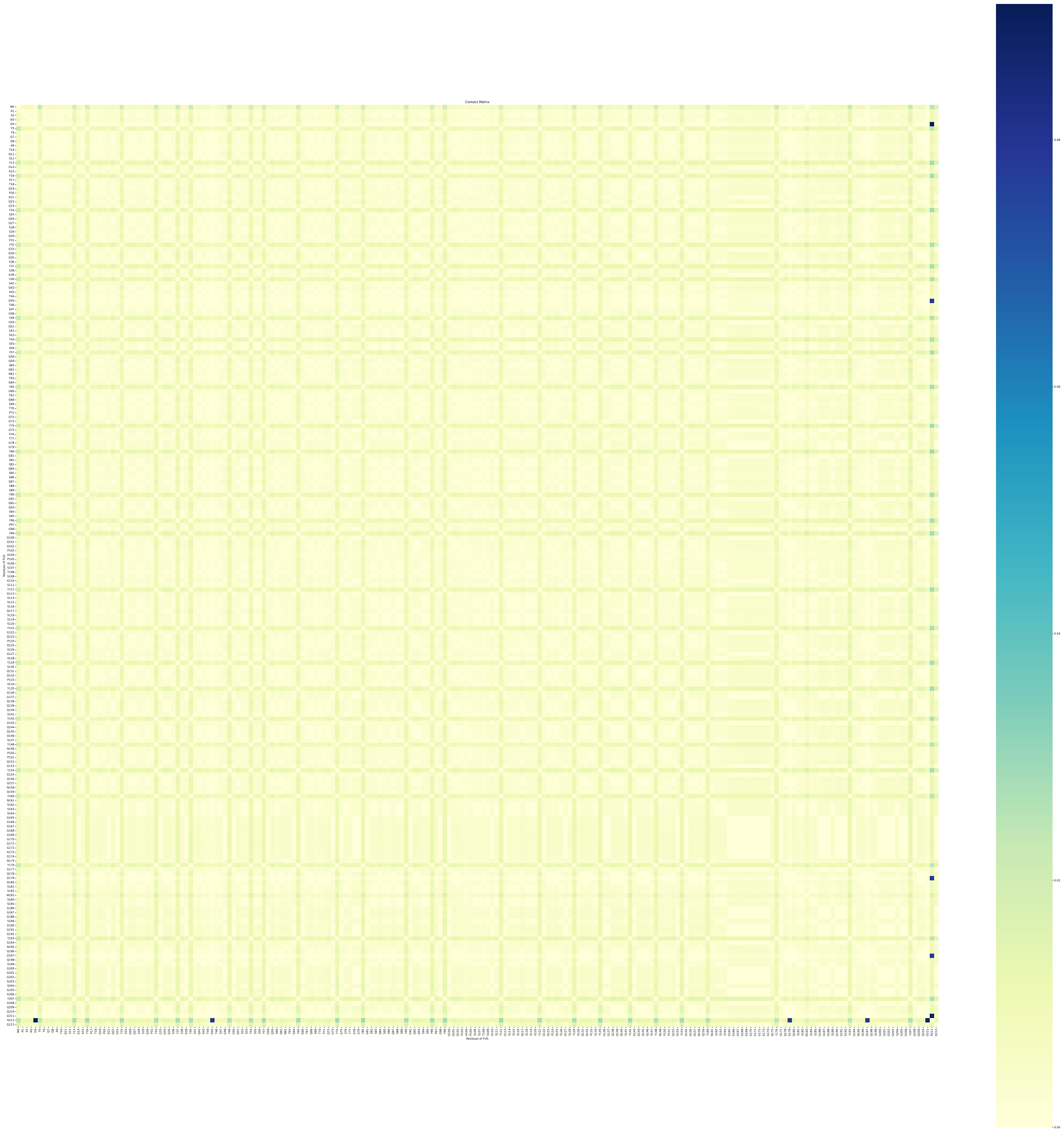

Figure S11: The probability of contact formation between residues of the FUS IDR in the bulk of its condensate.

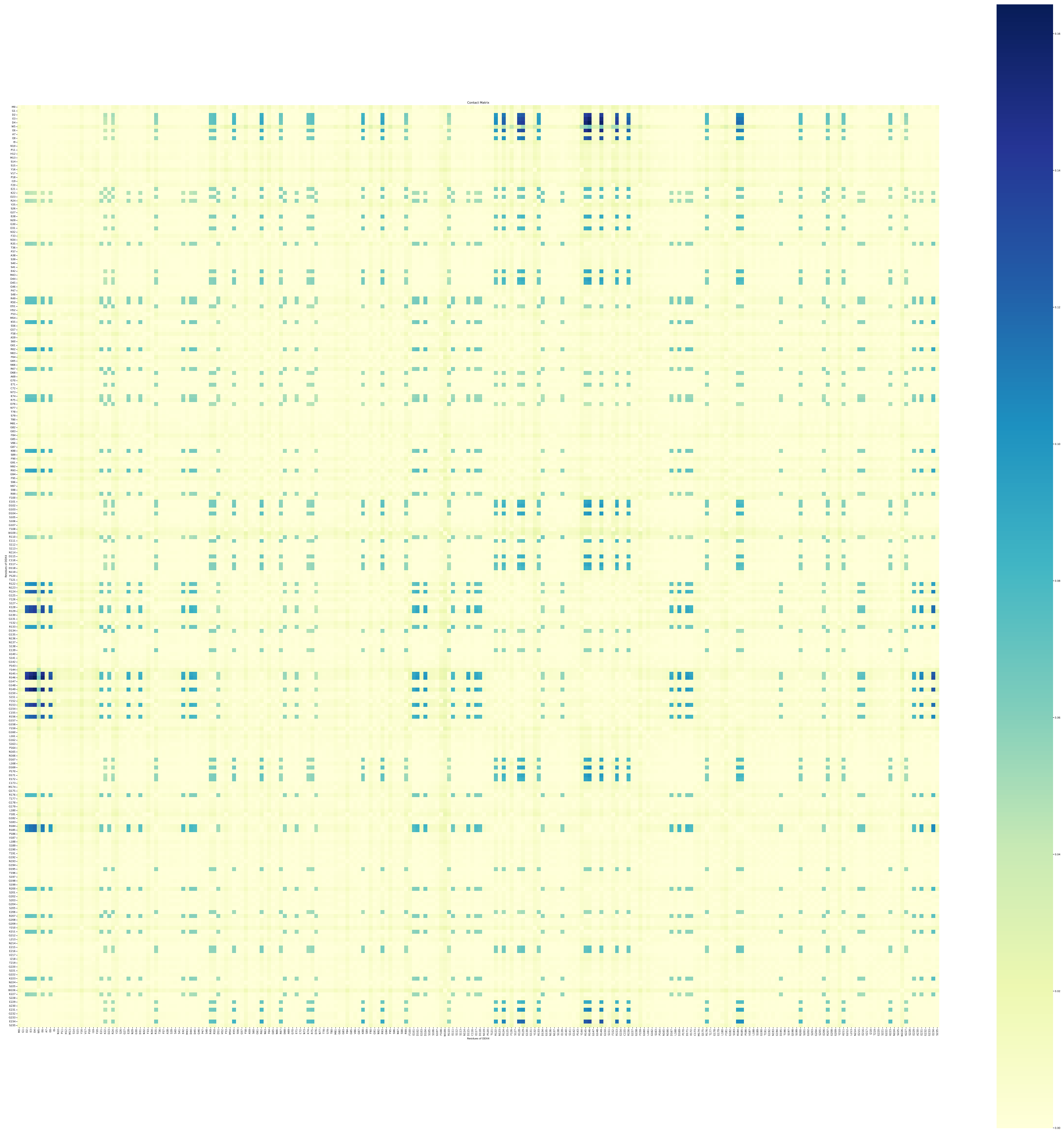

Figure S12: The probability of contact formation between residues of the DDX4 NTD IDR in the bulk of its condensate.

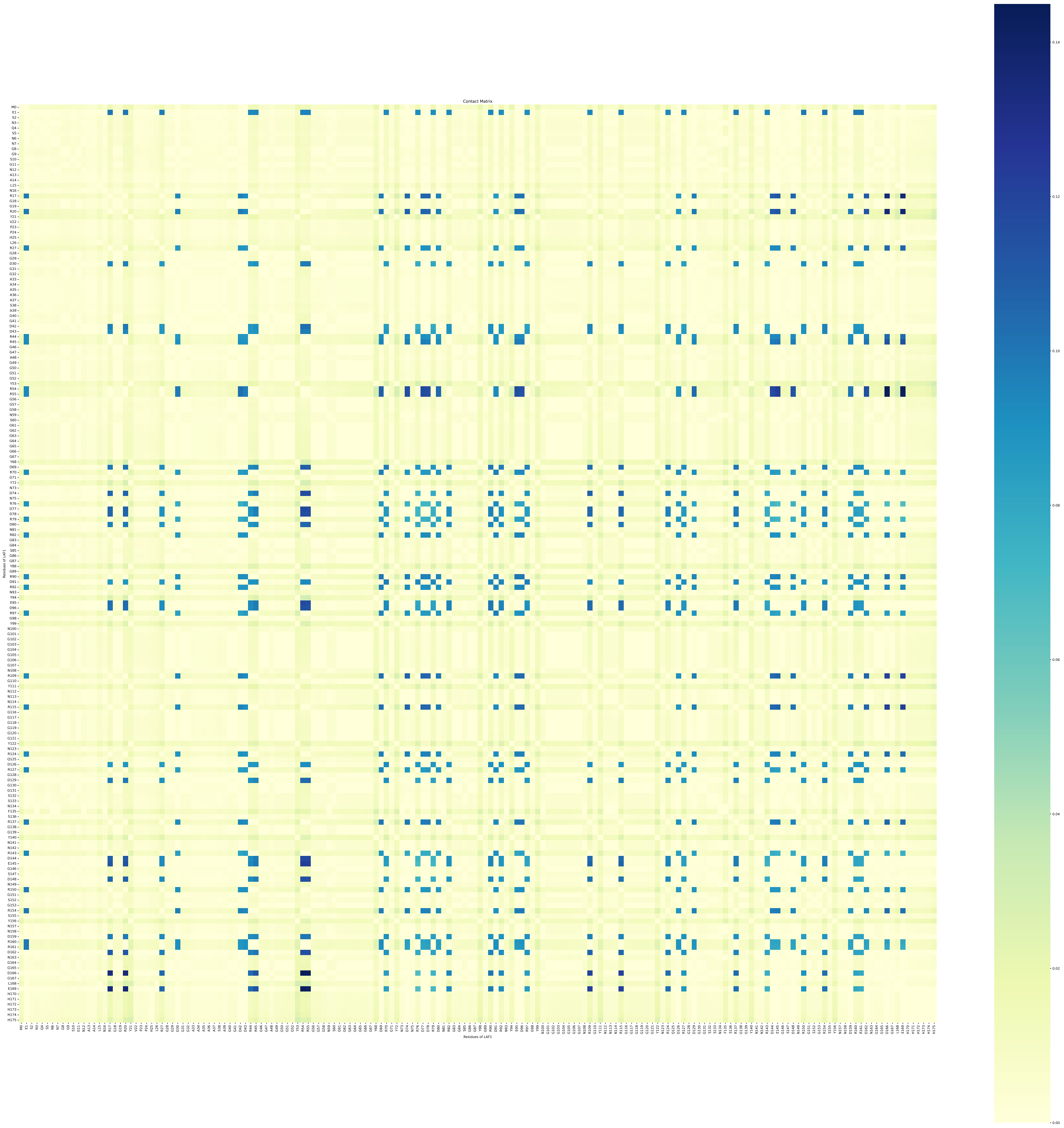

Figure S13: The probability of contact formation between residues of the LAF1-RGG IDR in the bulk of its condensate.

##### S4.1.1 Contact Blocks in IDR Chains

Domains in the IDR chains within which 'active' residues form contacts co-operatively were identified as contact 'blocks'. Fig. S14 shows the behavior of the joint-probabilities, the average size of the blocks, the number of blocks in an IDR chain, and the randomness of contact formation of the active residues within a block. The definition of a block was expanded based on the criterion of selecting active residues in the chain. The active residues are the ones that have a probability of making contact in the top $n^{th}$ percentile.

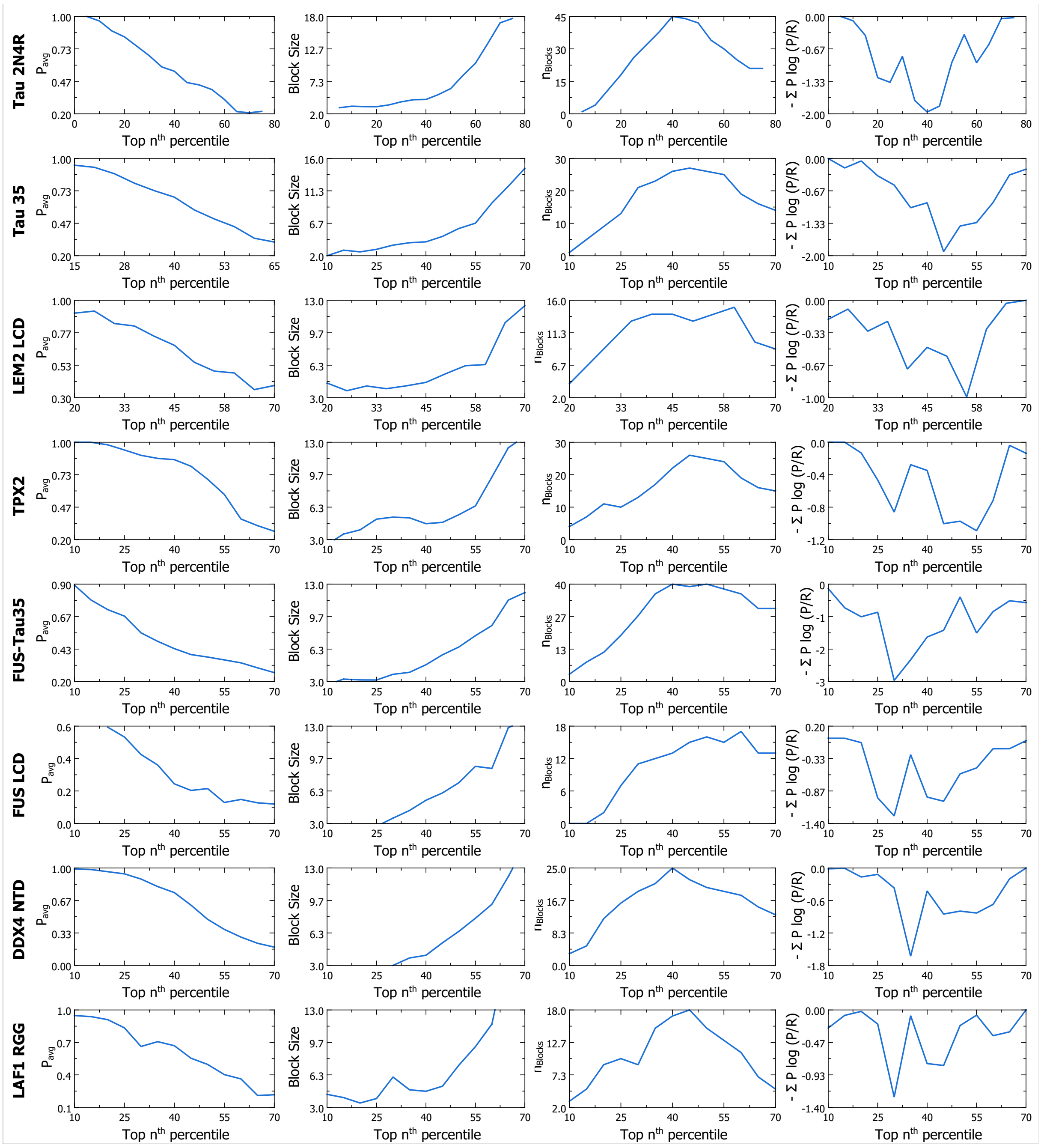


Figure S14: The properties of the block plotted against the criterion of selecting active residues in the chain. The average joint probabilities of a block, the average length of blocks, the number of the blocks, and the randomness of contact-formation of residues in blocks (measured by $-\sum \mathrm{P}\log(\mathrm{P}/\mathrm{R})$ ) are shown.

.

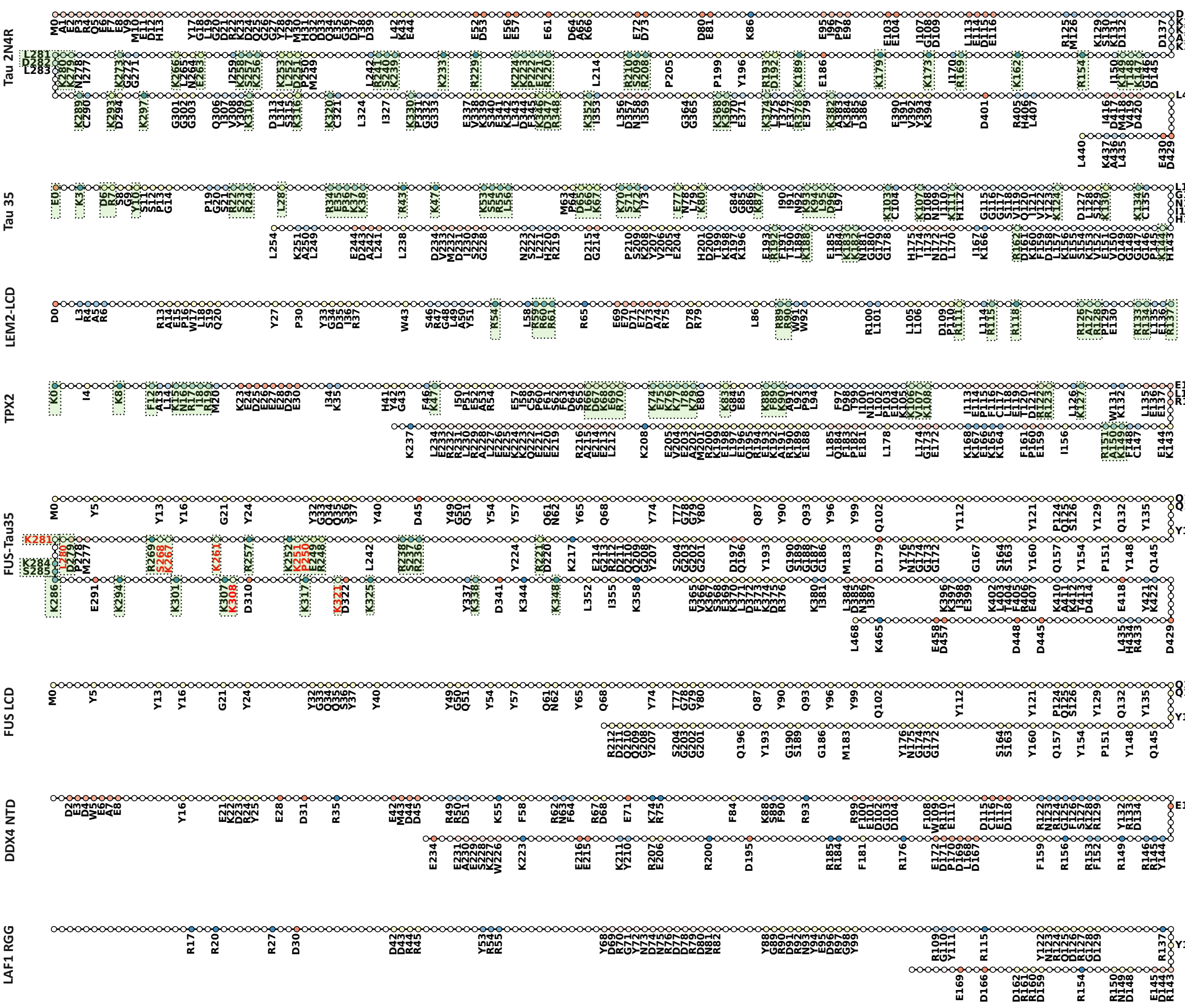


Figure S15: The contact-blocks of IDRs that are responsible for the stabilisation of the IDR condensates. The labeled residues represent the contact blocks in the IDR chain. The parts of the blocks that interact with the MT surface as highlighted in green. Residues labeled in red (FUS-Tau35) indicate the residues interacting with MT, but are not part of the blocks that stabilise the condensate bulk. Blocks with positive, negative, and neutral charges are coloured in blue, red, and yellow, respectively. The net charge per residue and the hydrophobicity of the blocks, and their neighborhood is shown in Fig. S16

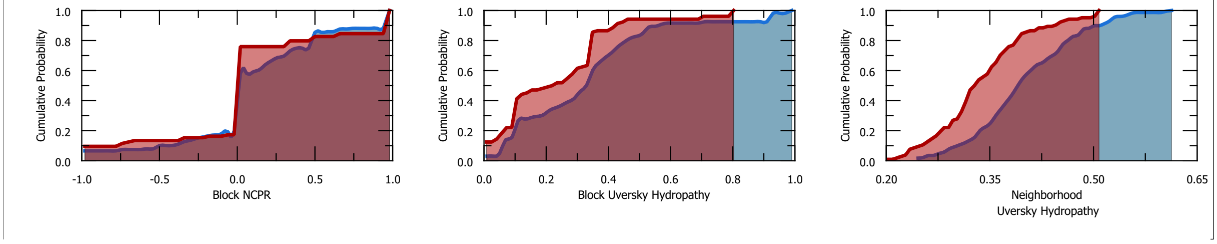


Figure S16: The cumulative distribution of charge and hydrophobicity of MT-wetting (blue) and non-wetting (red) IDRS. We observe that MT-wetting IDRs tend to have higher net-charge-per residue and uversky hydropathy in their contact blocks. We also observe a higher Uversky hydropathy in the neighborhood (±5 residues from the edge) of the contact blocks.

#### S4.2 MT-IDR Interactions

The contacts between IDR residues and $\alpha$- and $\beta$- tubulins were studied to understand the interactions that enable the wetting of the MT surface. Fig. S17 shows the most important interactions between IDRs and the tubulins, coloured according to the probability of interaction. Interactions are overwhelmingly found to be between the positive residues of the IDR and the negative residues of the MT surface.

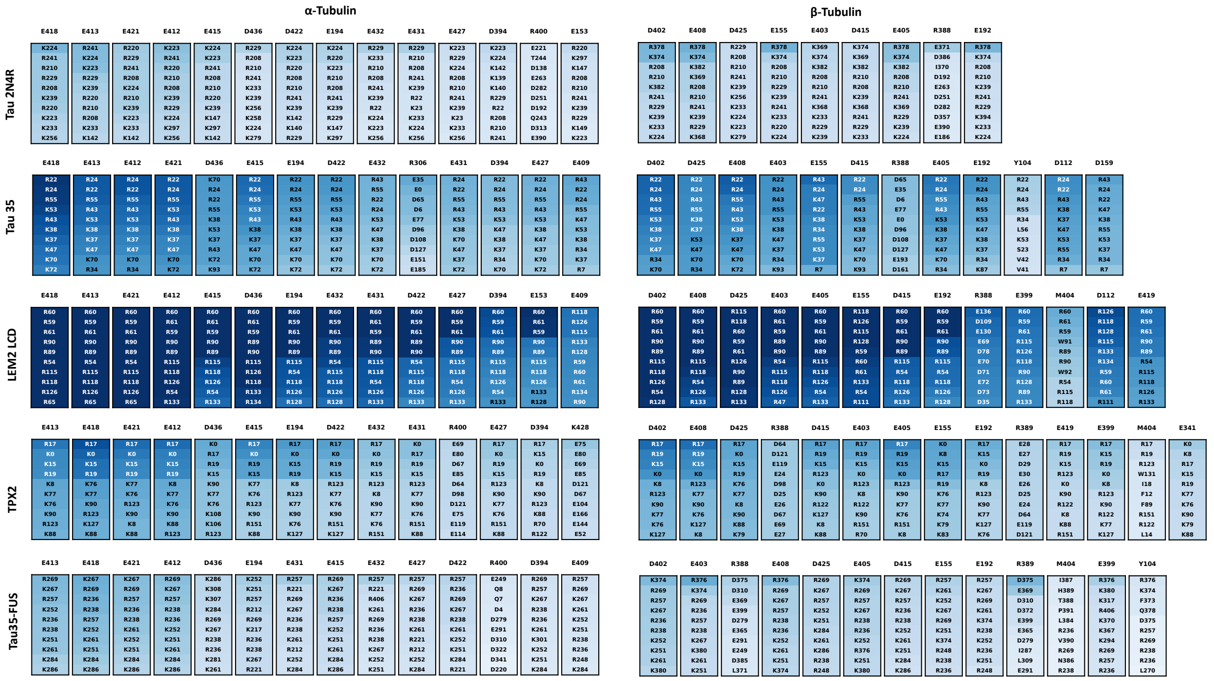


Figure S17: The most important contacts formed between the MT-surface and the residues of the IDR. Interactions with $\alpha$- and $\beta$- tubulins are shown separately. The residues numbers of all proteins are based on a 0-based index.

#### S4.3 Sequence Patterns of MT-wetting IDRs

A pattern of hydrophobic-positive domains was observed in the IDRs that wet the MT-surface. Our proposed model is also in agreement with the NARDINI analysis developed by Cohan et al. [S31]. The method involves the generation of an ensemble of scrambled sequences of a given composition, and subsequently setting up a composition-sepcific "null" model. Then, the z-score representing the deviation of patterns of the given sequence from the "null" model is computed. The deviation in z-scores of binary patterns of different residue types is shown in Fig. S18. A negative z-score indicates that the residue types are more "interlaced" between each other when compared to the null model. A positive z-score indicates that they are segregated from each other. An important distinction observed for the MT-wetting IDRs is the non-segregation of positive and hydrophobic residues. This agrees with our proposed model. Positive residues of IDR with hydrophobic residues in their vicinity are favored on the surface of the MT, enabling their condensates to wet the MT surface.

##### S4.3.1 Random-forest Model

To further investigate the factors that enable IDR residues to make contacts with the surface of the MT, a random forest regressor model was trained. The details of the models are described in this section.

The model was trained to predict the probability that a given residue would make any contact at the surface of the MT. Eight descriptors were used to train the model- two describing the residue itself (charge and Uversky hydropathy), and six descriptors describing the neighborhood of the residue (fraction of various residue types, namely, positive, negative, hydrophobic, aromatic, polar, and Glycine). The neighborhood is defined as residues in the sequence that are within $\pm 5$ residues from the central residue. The total dataset had 1543 data points (including residues of all the MT-wetting IDRs (Tau2N4R,

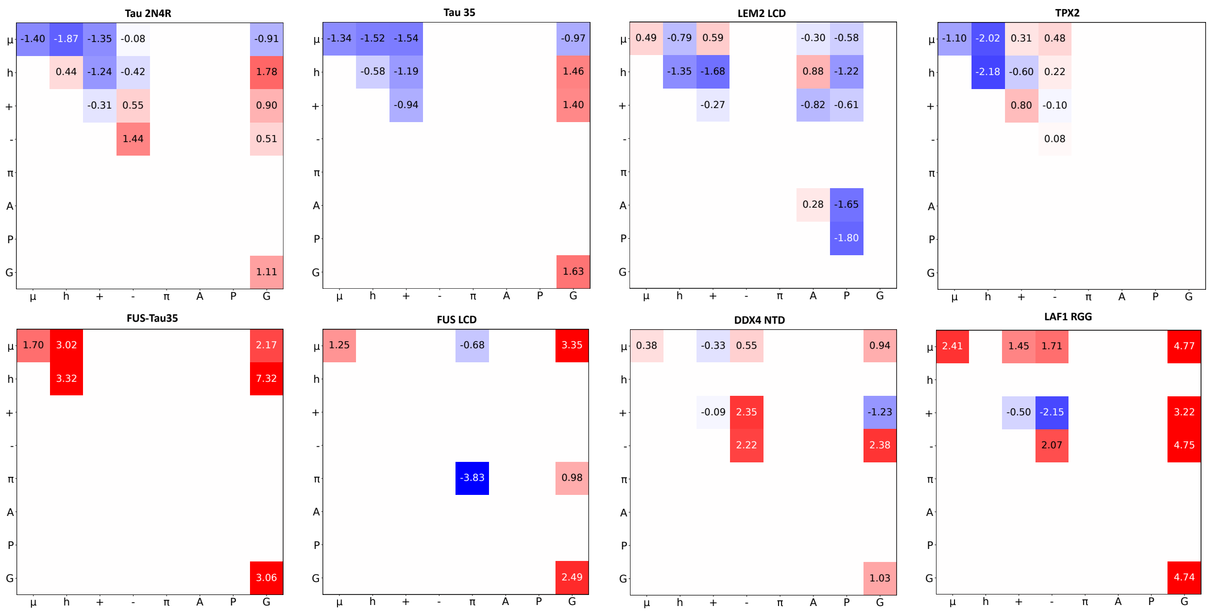


Figure S18: The NARDINI analysis to quantify binary patterns of residue types in a sequence. In general, it is observed that the residues that wet the surface of the MT have interlaced positive and hydrophobic residues.

Tau35, FUS-Tau35, LEM2 LCD, and TPX2). 20% of the data was not used for optimizing the hyperparameters or training the model.

Hyperparameter optimization was performed on the training set using a grid search procedure with five-fold cross-validation. The final model fitted on the training data was then evaluated on the held-out test data (see Fig. S19). To visualize and interpret the effect of individual descriptors, we computed and plotted partial dependence plots (PDPs) using the final trained model. The partial dependence plots provide an estimate of how the average MT-contact probability of residues vary with each descriptor in the dataset. We report the partial dependence of each descriptor for positive, negative, and uncharged residues separately (see Fig. S20-S22). Below, we discuss the dependence of MT-contact probabilities on their neighborhood in the IDR chain.

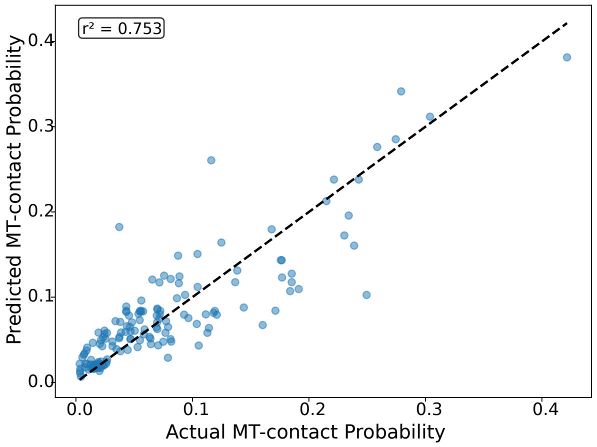


Figure S19: The actual MT-contact probabilities of IDR residues (not used while training the model) plotted against the values predicted by the random forest regression model.

For positive residues in the IDR chain, the presence of positive residues and the fraction of hydrophobic residues in the neighborhood is observed to have a positive effect on the mt-contact probability of the residue (see Fig. S20). This is again in agreement with our proposed model that IDRs with positive-hydrophobic domains enable IDR condensates to spread on the surface of MT. The presence of negative residues in the vicinity reduces the MT contact probabilities of positively charged residues. Presence of negative residues lowers the net-charge density close to positive residues, and therefore reduces the strength of positive-hydrophobic domains of the IDR with the negative-hydrophobic domains of the MT surface.

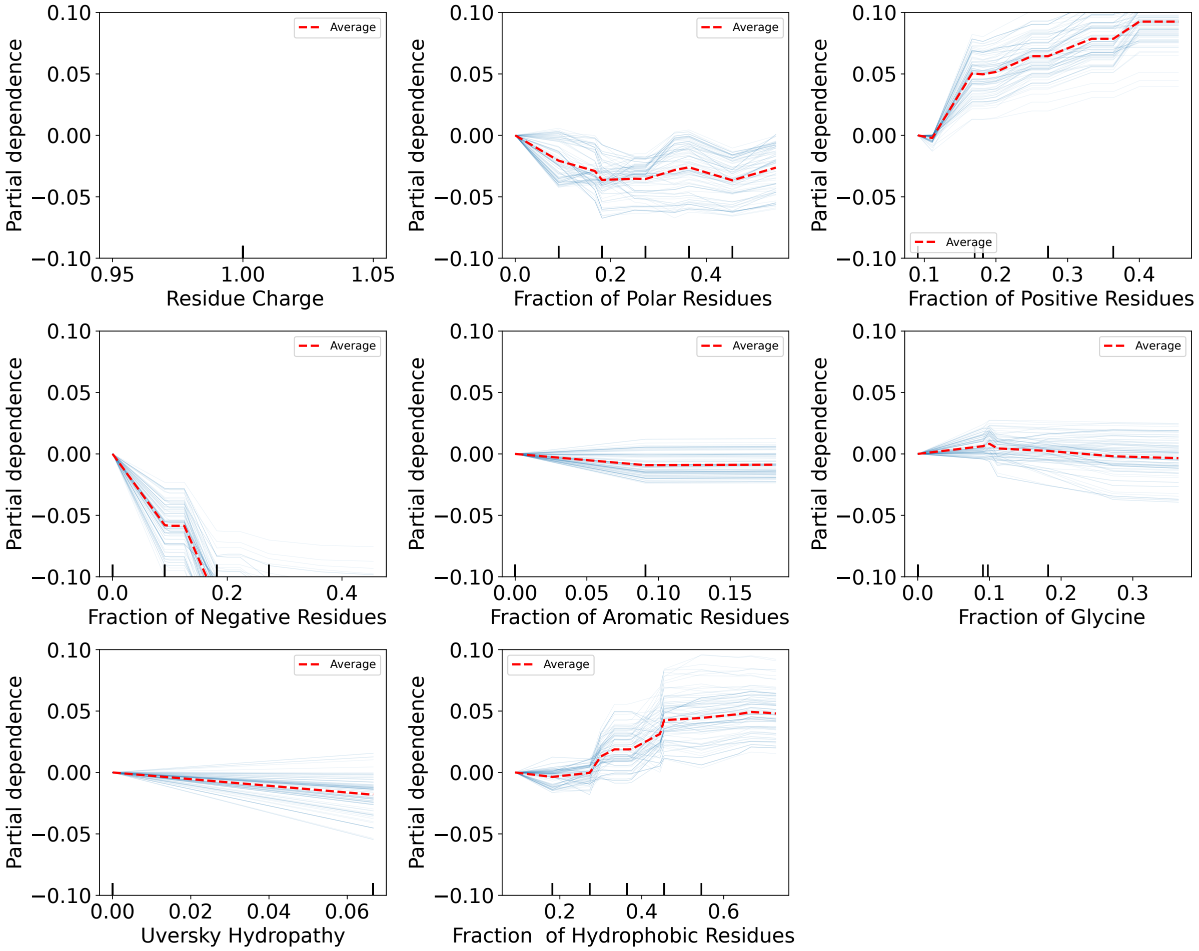


Figure S20: Partial dependence of MT-contact probability of positively charged IDR residues on their descriptors. The residue charge of positively charged residues is a constant. Therefore, the partial dependence plot for residue charge is left empty.

Negatively charged residues are often not preferred on the surface of the MT. Because of significantly lower hydrophobicity in the vicinity. However, the presence of positive charges in their vicinity can lead to a higher probability of binding on the surface. This is possibly because of the binding of the positive residues in the neighborhood onto the surface of the MT. This brings the negative residues close to the surface, where it can bind to the positively charged residues on the MT-surface.

For uncharged residues, the partial dependence plots suggest that the biggest factor that leads to an increase in contact probability is the presence of positive residues in the neighborhood. The positive charges bound to the MT surface bring them closer to the surface, enabling the formation of contacts at the surface.

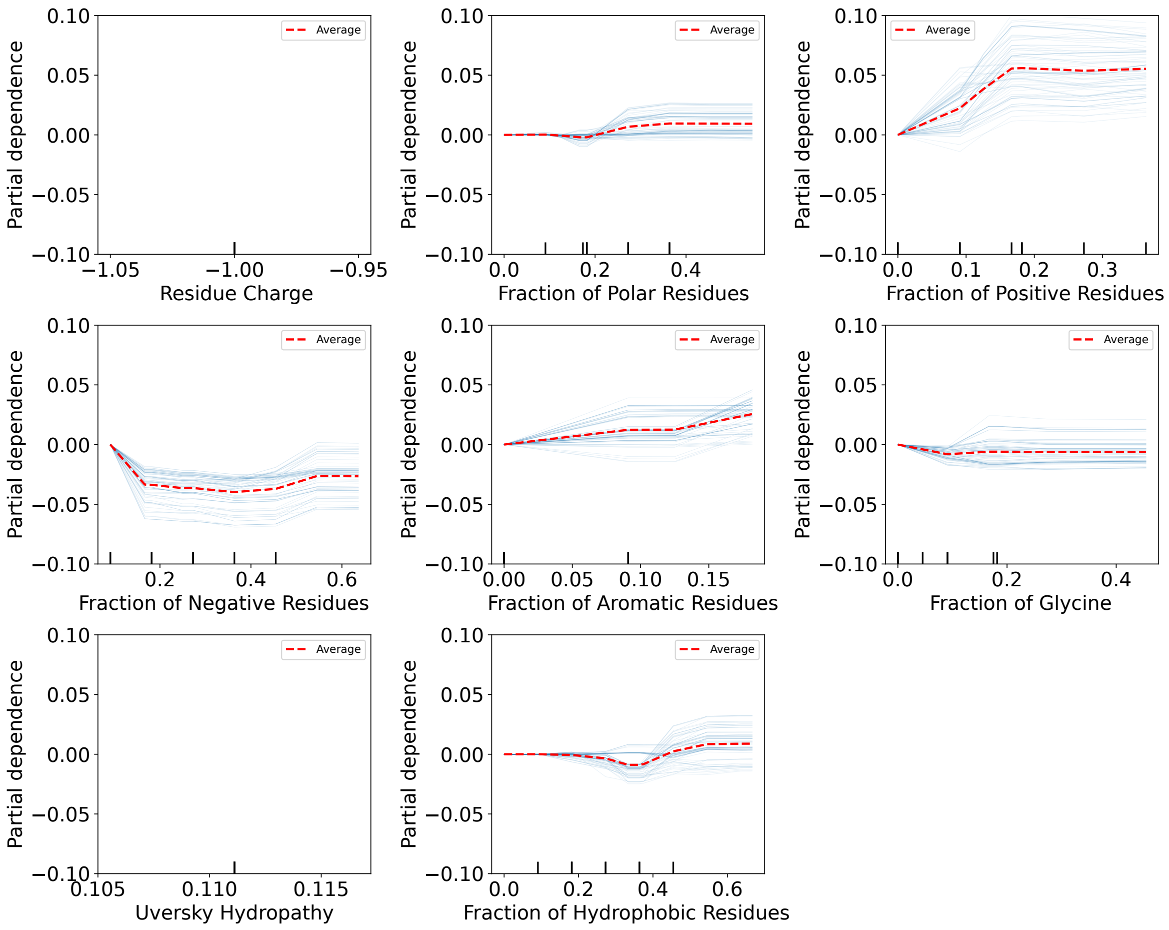


Figure S21: Partial dependence of MT-contact probability of negatively charged IDR residues on their descriptors. The residue charge and Uversky hydropathy are constant for negatively charged residues. Therefore, the partial dependence plots are left empty.

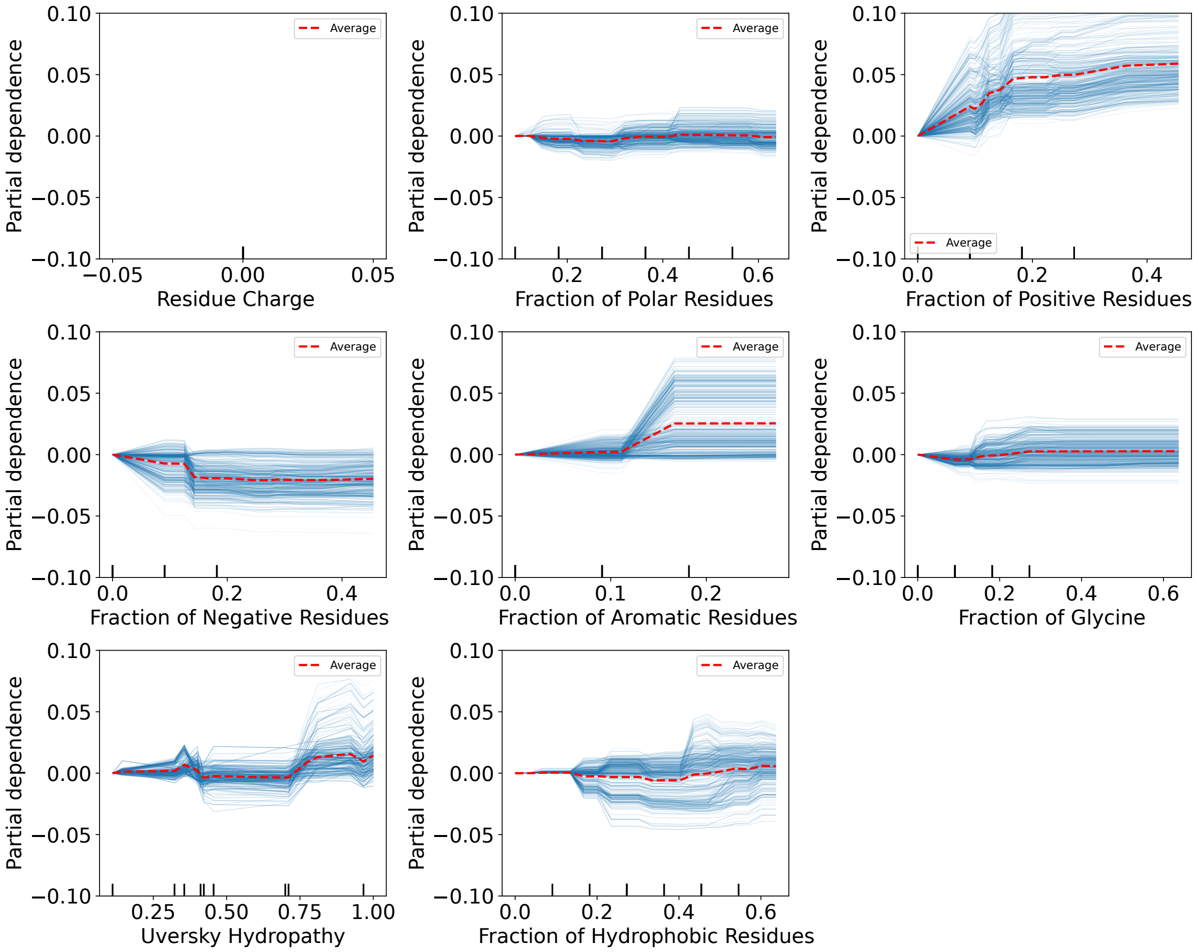


Figure S22: Partial dependence of MT-contact probability of non-charged IDR residues on their descriptors. The residue charge of positively charged residues is a constant. Therefore, the partial dependence plot for residue charge is left empty.

#### SI References


[S1] Alexander von Appen et al. “LEM2 Phase Separation Promotes ESCRT-Mediated Nuclear Envelope Reformation”. In: *Nature* 582.7810 (2020), pp. 115–118. DOI: 10.1038/s41586-020-2232-x.

[S2] John Jumper et al. “Highly Accurate Protein Structure Prediction with AlphaFold”. In: *Nature* 596.7873 (2021), pp. 583–589.

[S3] Mihaly Varadi et al. “AlphaFold Protein Structure Database in 2024: Providing Structure Coverage for over 214 Million Protein Sequences”. In: *Nucleic Acids Res.* 52.D1 (2024), pp. D368–D375.

[S4] Selina Wray et al. “Direct Analysis of Tau from PSP Brain Identifies New Phosphorylation Sites and a Major Fragment of N-Terminally Cleaved Tau Containing Four Microtubule-Binding Repeats”. In: *J. Neurochem.* 105.6 (2008), pp. 2343–2352.

[S5] Chen Lyu et al. “The Disease-Associated Tau35 Fragment Has an Increased Propensity to Aggregate Compared to Full-Length Tau”. In: *Front. Mol. Biosci.* 8 (2021), p. 779240. DOI: 10.3389/fmolb.2021.779240.

[S6] Sunhwan Jo et al. “CHARMM-GUI PDB Manipulator for Advanced Modeling and Simulations of Proteins Containing Nonstandard Residues”. In: *Adv. Protein Chem. Struct. Biol.* 96 (2014), pp. 235–265.

[S7] Sang-Jun Park et al. “CHARMM-GUI PDB Manipulator: Various PDB Structural Modifications for Biomolecular Modeling and Simulation”. In: *J. Mol. Biol.* 435.14 (2023), p. 167995.

[S8] Alexander W Bird and Anthony A Hyman. “Building a Spindle of the Correct Length in Human Cells Requires the Interaction between TPX2 and Aurora A”. In: *J. Cell Biol.* 182.2 (2008), pp. 289–300. DOI: 10.1083/jcb.200802005.

[S9] David K Moss, Andrew Wilde, and Jon D Lane. “Dynamic Release of Nuclear RanGTP Triggers TPX2-Dependent Microtubule Assembly During the Apoptotic Execution Phase”. In: *J. Cell Sci.* 122.5 (2009), pp. 644–655. DOI: 10.1242/jcs.037259.

[S10] Xuli Qi et al. “UHRF1 Promotes Spindle Assembly and Chromosome Congression by Catalyzing EG5 Polyubiquitination”. In: *J. Cell Biol.* 222.11 (2023), e202210093. DOI: 10.1083/jcb.202210093.

[S11] Raymundo Alfaro-Aco, Akanksha Thawani, and Sabine Petry. “Structural Analysis of the Role of TPX2 in Branching Microtubule Nucleation”. In: *J. Cell Biol.* 216.4 (2017), pp. 983–997. DOI: 10.1083/jcb.201607060.

[S12] Raymundo Alfaro-Aco and Sabine Petry. “How TPX2 Helps Microtubules Branch Out”. In: *Cell Cycle* 16.17 (2017), pp. 1560–1561.

[S13] Mina Farag et al. “Phase Separation of Protein Mixtures Is Driven by the Interplay of Homotypic and Heterotypic Interactions”. In: *Nat. Commun.* 14 (2023), p. 5527.

[S14] Chih-Chia Chang and Scott M Coyle. “Regulatable Assembly of Synthetic Microtubule Architectures Using Engineered Microtubule-Associated Protein-IDR Condensates”. In: *J. Biol. Chem.* 300.8 (2024). DOI: 10.1016/j.jbc.2024.107544.

[S15] Benjamin S Schuster et al. “Identifying Sequence Perturbations to an Intrinsically Disordered Protein that Determine its Phase-Separation Behavior”. In: *Proc. Natl. Acad. Sci. USA* 117.21 (2020), pp. 11421–11431.

[S16] Rahul K Das and Rohit V Pappu. “Conformations of Intrinsically Disordered Proteins Are Influenced by Linear Sequence Distributions of Oppositely Charged Residues”. In: *Proc. Natl. Acad. Sci. USA* 110.33 (2013), pp. 13392–13397.

[S17] Alex S Holehouse et al. “CIDER: Resources to Analyze Sequence-Ensemble Relationships of Intrinsically Disordered Proteins”. In: *Biophys. J.* 112.1 (2017), pp. 16–21.

[S18] Vladimir N Uversky. “Natively Unfolded Proteins: a Point Where Biology Waits for Physics”. In: *Protein Sci.* 11.4 (2002), pp. 739–756.

[S19] Erik W Martin et al. “Sequence Determinants of the Conformational Properties of an Intrinsically Disordered Protein Prior to and Upon Multisite Phosphorylation”. In: *J. Am. Chem. Soc.* 138.47 (2016), pp. 15323–15335.

[S20] Tong Guo, Wendy Noble, and Diane P Hanger. “Roles of Tau Protein in Health and Disease”. In: *Acta Neuropathol.* 133 (2017), pp. 665–704.

[S21] Franziska Decker et al. “Autocatalytic Microtubule Nucleation Determines the Size and Mass of Xenopus Laevis Egg Extract Spindles”. In: *Elife* 7 (2018), e31149.

[S22] Gregory L Dignon et al. “Sequence Determinants of Protein Phase Behavior from a Coarse-Grained Model”. In: *PLOS Comput. Biol.* 14.1 (2018), e1005941. DOI: 10.1371/journal.pcbi.1005941.

[S23] Felipe J Blas et al. “Vapor-Liquid Interfacial Properties of Fully Flexible Lennard-Jones Chains”. In: *J. Chem. Phys.* 129.14 (2008). DOI: 10.1063/1.2989115.

[S24] Kevin S Silmore, Michael P Howard, and Athanassios Z Panagiotopoulos. “Vapour–Liquid Phase Equilibrium and Surface Tension of Fully Flexible Lennard–Jones Chains”. In: *Mol. Phys.* 115.3 (2017), pp. 320–327. DOI: 10.1080/00268976.2016.1262075.

[S25] Giulio Tesei et al. “Accurate Model of Liquid–Liquid Phase Behavior of Intrinsically Disordered Proteins from Optimization of Single-Chain Properties”. In: *Proc. Natl. Acad. Sci. USA* 118.44 (2021), e2111696118. DOI: 10.1073/pnas.2111696118.

[S26] John Shipley Rowlinson and Benjamin Widom. *Molecular Theory of Capillarity*. Courier Corporation, 2013. DOI: 10.1002/bbpc.19840880621.

[S27] Fan Cao et al. “A Coarse-grained Model for Disordered and Multi-Domain Proteins”. In: *Protein Sci.* 33.11 (2024), e5172. DOI: 10.1002/pro.5172.

[S28] Ana B Asenjo et al. “Structural Model for Tubulin Recognition and Deformation by Kinesin-13 Microtubule Depolymerases”. In: *Cell Reports* 3.3 (2013), pp. 759–768.

[S29] Sunhwan Jo et al. “CHARMM-GUI: A Web-Based Graphical User Interface for CHARMM”. In: *J. Comput. Chem.* 29.11 (2008), pp. 1859–1865.

[S30] Jesús A. Izaguirre, Chris R. Sweet, and Vijay S. Pande. “Multiscale Dynamics of Macromolecules Using Normal Mode Langevin”. In: *Pac. Symp. Biocomput.* 15 (2010), pp. 240–251. DOI: 10.1142/9789814295291_0026.

[S31] Megan C Cohan et al. “Uncovering Non-Random Binary Patterns Within Sequences of Intrinsically Disordered Proteins”. In: *J. Mol. Biol.* 434.2 (2022), p. 167373. DOI: 10.1016/j.jmb.2021.167373.
